# Fundamental trade-offs in the robustness of biological systems with feedback regulation

**DOI:** 10.1101/2024.09.25.614654

**Authors:** Nguyen Hoai Nam Tran, An Nguyen, Tasfia Wasima Rahman, Ania-Ariadna Baetica

## Abstract

Natural biological systems use a complex network of feedback regulation to effectively respond to their changing environment. Even though in engineered systems we understand how accurate feedback can be depending on the electronic or mechanical parts that it is implemented with, we largely lack a similar theoretical framework to study biological feedback. Specifically, it is not fully understood or quantified how accurate or robust the implementation of biological feedback actually is. In this paper, we study the sensitivity of biological feedback to variations in biochemical parameters using five example circuits: positive autoregulation, negative autoregulation, double-positive feedback, positive-negative feedback and double-negative feedback (the toggle switch). We find that of these examples of biological feedback are subjected to fundamental trade-offs, and we propose multi-objective optimisation as a framework to study them. The impact of this work is to improve robust circuit design for synthetic biology and to improve our understanding of systems biology.

## 1 INTRODUCTION

In biological systems, feedback can regulates and maintains the stability of the internal environments of organisms and cells, ensuring homeostasis despite changes in their external environments (*1 –3*). Additionally, feedback in biological systems can generate dynamic responses such as oscillations, signal amplification, and facilitate rapid switching between multiple states (*4 –8*). These roles of biological feedback are crucial for processes such as gene expression, metabolic regulation, and adaptation to environmental changes. Despite the prevalence of feedback in natural biological systems, its properties are not fully understood and moreover, its similarities and differences with engineered feedback remain understudied (*9, 10*). In this work, we study the robustness of biological feedback to variations in its biochemical implementation and its system design properties.

In mechanical and electrical systems, feedback can be used to improve a system’s robustness against disturbances (*11*). For example, negative feedback in electronic circuits can improve robustness for a specific input range though it can also reduce overall gain and introduce fragility elsewhere. Indeed, it is well-known in engineering that feedback cannot provide complete robustness against all disturbances and that the use of feedback can introduce fundamental limitations on the system’s performance (*12, 13*). The use of feedback in engineering can create performance trade-offs in terms of speed, stability, and accuracy that must be carefully balanced during system design.

In contrast, it remains mostly unknown whether feedback introduces similar trade-offs in biological systems (*14*). Previous research on this topic has experimentally validated a trade-off between efficiency and robustness in the glycolysis pathway of *Saccharomyces cerevisiae* (*15*). Other trade-offs were uncovered through modelling and mathematical analysis (*16 –20*). However, we are still missing a general framework to study biological feedback and its trade-offs.

A second key difference between feedback in engineered versus biological systems is that inside cells, feedback is implemented with biochemical reaction rates that can vary across many orders of magnitude and that can change due to mutations, and internal and external stimuli (*21*). Even in synthetically engineered biological circuits, the biochemistry of a feedback loop can be altered by its genetic, cellular, and extracellular contexts (*22, 23*). Largely, it remains unclear how robust biological feedback is to variations in the biochemical reaction rates that implement it (*10, 24*).

In this paper, we quantify the robustness of five well-known models of biological motifs to variations in their biochemical parameters using sensitivity analysis. Our motivation for selecting these five motifs is that they represent simple synthetic circuits with feedback regulation that have been both modelled and experimentally constructed. First, we consider the mathematical models of positive and negative autoregulation (*1*), the toggle switch (*25*), the double-positive feedback motif (*26*), and the positive-negative feedback motif (*27*). We construct sensitivity functions that capture how each motif’s output changes with variations in its biochemical parameters. Subsequently, we pose and solve the multiobjective optimization problems of simultaneously minimizing pairs of sensitivity functions to determine the Pareto-optimal implementations of each motif.

We find that three of these biological examples are robust to variations in their biochemical parameters, while the negative autoregulation motif and the positive-negative feedback motif are constrained by robustness trade-offs. These results offer insight into how to optimally design robust, reliable synthetic circuits that can function robustly in changing environments.

The impact of our research is in providing a framework to study how robustness is allocated feedback in biological systems and what trade-offs are created by the use of biological feedback. Additionally, our research enhances our knowledge of systems biology and informs the system design of synthetic biological circuits.

## 2 RESULTS AND DISCUSSION

### 2.1 Sensitivity Analysis

Intracellular feedback loops depend on physical parameters, such as the cooperativity and the binding affinity of biomolecules. Understanding how changes in these physical parameters change the concentrations of biochemical species in a system will give insight into the properties of feedback loops. Sensitivity analysis quantifies this relationship, revealing which parameters have the greatest impact on the behaviour of a biological system (*2, 28*).

Throughout this paper, we will use sensitivity analysis to assess the robustness of biological systems with feedback to changes in their biochemical parameters such as binding and unbinding rates, cooperativities, and degradation rates. Here, we introduce the sensitivity function that we will use throughout the paper to determine how sensitive feedback loops are to variations in these parameters.

We assume that the biological systems we consider can be modelled using a system of nonlinear ordinary differential equations. The model is of the form:

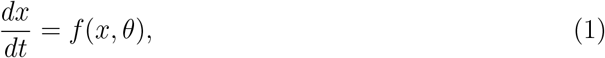

where *x* = (*x*^1^, …, *x*^*n*^) is a vector of *n* biochemical species, *θ* = (*θ*^1^, …, *θ*^*k*^) is a vector of *k* biochemical parameters, and *f* is a nonlinear, continuous function representing the dynamics of *x*. Assuming that a steady-state exists, we must have that

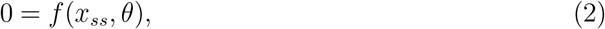

where *x*_*ss*_ is the vector of biochemical species *x* evaluated at steady-state. The steady-state condition creates an implicit equation with variables *x*_*ss*_ and *θ*.

A simple measure of the sensitivity of biological systems to their biochemical parameters is given by the sensitivity function at steady-state. For a biochemical species 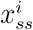 at steady-state, its sensitivity with respect to the parameter *θ*_*j*_ is given by

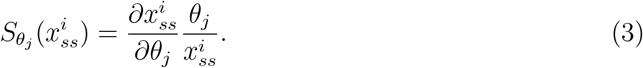

Here, 1 ≤ *i* ≤ *n* and 1 ≤ *j* ≤ *k*. The sensitivity function can be thought of as a ratio between the fractional change in the biochemical species 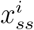, given by 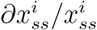, to the fractional change in its dependent parameter *θ*_*j*_. If 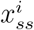 represents a biochemical species dependent on a parameter *θ*_*j*_, then 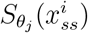 represents the ratio of the fractional change in the species to the fraction that we have perturbed the biochemical parameter from its original value by. To extend this idea further, the sensitivity function of a set of biochemical species at steady-state, 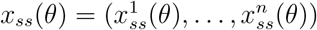 with respect to *θ* is simply

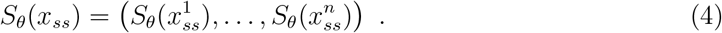

We briefly note that there are other methods to quantify the sensitivity of a system to variations in its parameters (*12, 28*), but that we choose this local derivative-based sensitivity function for its simplicity and ease of application to nonlinear biological systems.

### 2.2 Pareto-Optimality

Throughout this paper, we will use the multi-objective optimization (MOO) theoretical framework to simultaneously minimize multiple sensitivity functions. This will ensure robustness in our biological systems to variations in multiple of their biochemical parameters. Multi-objective optimization (MOO) is a type of optimization problem in which more than one objective function is optimized at the same time. In MOO problems, the objectives can conflict with each other, meaning that improving one objective may lead to a deterioration in another. The goal of MOO problems is not to find a single “best” solution but rather a set of optimal solutions, known as Pareto-optimal, where no single objective can be improved without worsening at least one other objective. This set of optimal solutions is called the Pareto front.

We formulate the multi-objective optimization problem for our examples as follows:

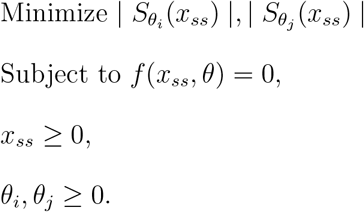

We assume that *x*_*ss*_ is the set of biochemical species at steady-state and that *θ*_*i*_ and *θ*_*j*_ are two of the biochemical parameters in the model in Equations (1) and (2).With this formulation, we aim to find the Pareto-optimal parameter set that simultaneously minimises a pair of sensitivity functions.

We note that MOO problems have previously been solved for the system design of synthetic circuits (*29*) and to understand other biological systems in (*30 –32*).

### 2.3 Biological Systems and their Robustness Trade-offs

#### 2.3.1 Positive autoregulation is unconstrained by trade-offs

Positive autoregulation occurs when a gene or protein enhances its own production by stimulating its own expression (*1*). In this type of regulatory feedback loop, the product of the gene, such as a protein or RNA, increases the rate of its own synthesis, leading to an amplification of its expression. This mechanism helps sustain high levels of gene or protein expression, which is crucial for processes like cell differentiation, developmental pathways, and signal amplification in biological systems.

The non-dimensionalised model of positive autoregulation from (*1*) is given by the following ordinary differential equation in species *x*:

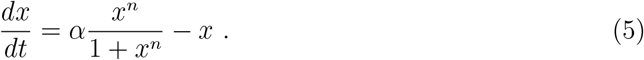

In this model, *n* represents the cooperativity constant of species *x*. Cooperativity is a molecular phenomenon where the binding of a molecule such as a ligand to a protein or enzyme influences the binding of additional molecules, either by enhancing or reducing more binding. Parameter *α* represents the ratio of the production rate to the degradation rate per unit of dissociation constant concentration. Throughout the paper, we refer to *α* as the “feedback strength”. Both parameters *α* and *n* are unitless quantities. We illustrate this biological circuit’s diagram in Figure 2A).

**Figure 1:**
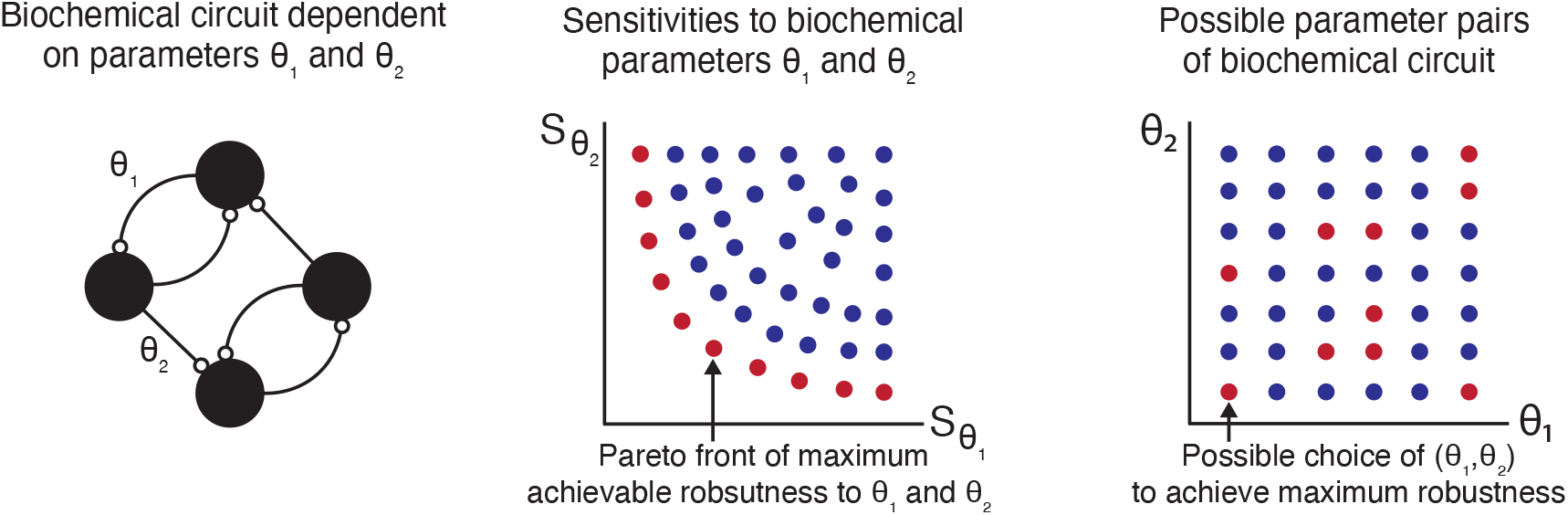
Biochemical circuits depend on physical parameters. Minimizing the sensitivity to these physical parameters allows us to find the most robust circuit designs.

**Figure 2:**
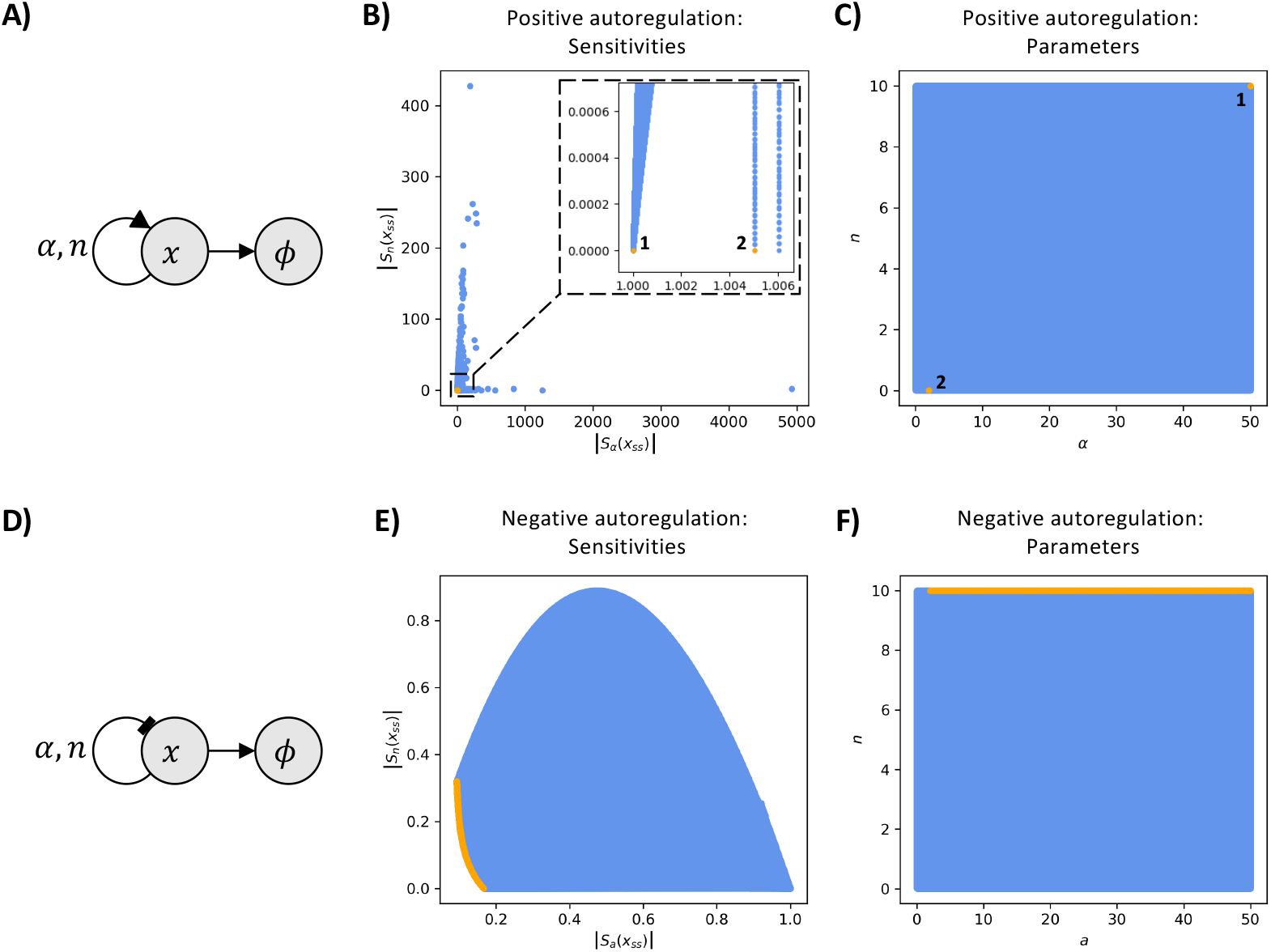
Pareto fronts of the sensitivity functions for the positive and negative autoregulation systems. **Panels A) and D)** show biological diagrams of the positive (A) and the negative (C) autoregulation of a biochemical species *x*. In **panel A)**, the pointed arrow represents positive feedback, where *x* enhances its own production, while in **panel C)**, the blunt arrow depicts negative feedback, where *x* represses its own production. We assume that species *x* also undergoes degradation and dilution within the cell, as indicated by the arrows pointed to the empty set symbol. The arrows indicate the parameter dependencies: *n* represents the binding cooperativity, while *α* represents the ratio of the production rate to the degradation rate per unit of dissociation constant concentration. We refer to *α* throughout the paper as the “feedback strength”. **Panels B) and E)** illustrate the Pareto fronts (orange) corresponding to the simultaneous minimization of the absolute values of the sensitivities with respect to the feedback strength, *α*, and to the cooperativity, *n* (blue). In **panel B)**, the Pareto front consists of two points, labelled 1 and 2. However, point 2 is identified as a numerical artifact of the sampling, meaning the true Pareto front contains only point 1. Solving the MOO problem shows that there is no trade-off between its two sensitivities. In **panel E)**, the Pareto front for the negative autoregulation system is represented by the orange curve. Solving the MOO problem shows that there is a trade-off between the sensitivities of the negative autoregulation circuit. **Panels C) and F)** illustrate the corresponding sampled parameters (blue) and the parameters responsible for the Pareto fronts (orange). In **panel C)**, two Pareto-optimal parameter pairs are identified for the MOO problem of positive autoregulation. However, since point 2 is a numerical artifact, only point 1 at maximal feedback strength and maximal cooperativity contributes to the Pareto front. In **panel F)** the Pareto front is shown to be created by parameters with maximal cooperativity and feedback strength values of 2 or greater.

The steady-states of Equation (5) occur when

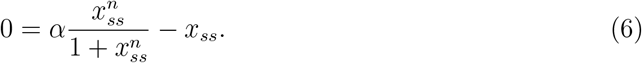

We note that *x*_*ss*_ = 0 is always a solution to Equation (6). However, the sensitivity function *S*_*θ*_(*x*_*ss*_) is ill-defined when *x*_*ss*_ = 0 due to division by zero. Hence the zero solution must be removed for our analysis. Thus, Equation (6) reduces to

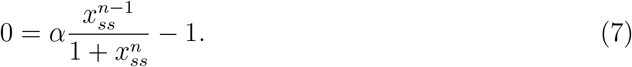

Equation (7) can have up to two solutions, depending on the choice of parameters. When one solution exists, it is always stable. When two solutions exist, one is stable and the other is unstable. We are always interested in the stable solution, as this is the one that can be observed and measured.

In the Supplementary Materials, we provide the derivations of the relative sensitivities with respect to biochemical parameters *α* and *n*. The resulting sensitivity functions are

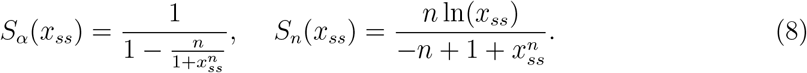

We note that sensitivity *S*_*α*_(*x*_*ss*_) ≥ 1 from Equation (8). We solve the MOO problem of simultaneously minimizing the absolute values of these sensitivities and illustrate the result in Figure 2B). We observe that there is no trade-off between the two sensitivities for the positive autoregulation circuit. Intuitively, it makes sense that positive feedback is not sensitive to the parameters in its implementation due to its ability to amplify signals to steady-state. This amplification may be insensitive to the biochemical parameter values. With these properties, positive autoregulation can make clear and decisive responses that are particularly useful in biological processes that require an all-or-nothing decision.

#### 2.3.2 Negative autoregulation is constrained by a fundamental trade-off

Negative autoregulation is a regulatory mechanism in biological systems where a gene or protein inhibits its own expression (*1*). In this process, the product of a gene represses its own transcription, effectively reducing the level of its production. The role of negative autoregulation in biological systems is to quickly respond to changes in environmental conditions or cellular needs.

Previous research on the robustness of negative intracellular feedback has found that it is robust to variation in the cooperativity of the biochemical species, but sensitive to its production and degradation rates at specific values of these rates (*24*). We extend this result to reveal that a trade-off constrains negative autoregulation. We find that negative feedback cannot be robust to both variations in both cooperativity and feedback strength.

We start by considering a system of one biochemical species undergoing negative autoregulation. We assume the species is produced via Hill kinetics with cooperativity constant *n* and that its feedback strength is *α* (*1*). This system is illustrated in Figure 2C) and modelled by the following non-dimensionalized differential equation model:

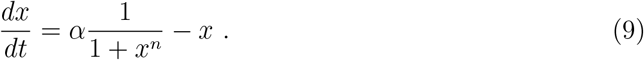

The steady state of this system occurs when

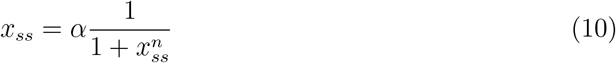

We note that Equation (10) has a unique steady-state solution. We want to assess the sensitivity of this steady state to changes in each of its two dependent parameters: *α* and *n*. We will therefore inspect the following sensitivity functions:

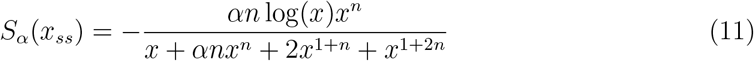

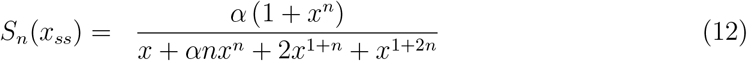

In Figure 2D), we note that there is a trade-off between the sensitivity to cooperativity and the sensitivity to feedback strength. This implies that the negative autoregulation system must have either some sensitivity to changes in feedback strength or to changes in cooperativity. Therefore, for a negative autoregulation system with cooperativity *n*, a higher value of the feedback strength will provide robustness to small changes in the feedback strength, and vice versa. To build a negative autoregulation circuit that is highly robust to changes in parameters, our modelling suggests that the feedback strength *α* should be greater than value two and the cooperativity *n* should be as high as possible. Clearly, finding the trade-off between the sensitivities provides valuable information about the system design of a robust negative autoregulation synthetic circuit.

#### 2.3.3 The double-positive feedback circuit is unconstrained by trade-offs

A biological system with two positive feedback loops, such as those found in T-cell differentiation, can reinforce a precursor cell’s commitment to a specific lineage (*33*). These two feedback loops ensure a robust and irreversible decision, maintaining the differentiated state even after initial signals fade, and preventing partial differentiation. This design can be used in synthetic biology for stable state transitions or decision-making processes in cells (*34*).

The double-positive feedback loop, illustrated in Figure 3A, consists of two mutually promoting species, *x* and *y*, that each undergo degradation. The non-dimensionalized model of the double-feedback system can be described by the two ODEs in Equation (13).

**Figure 3:**
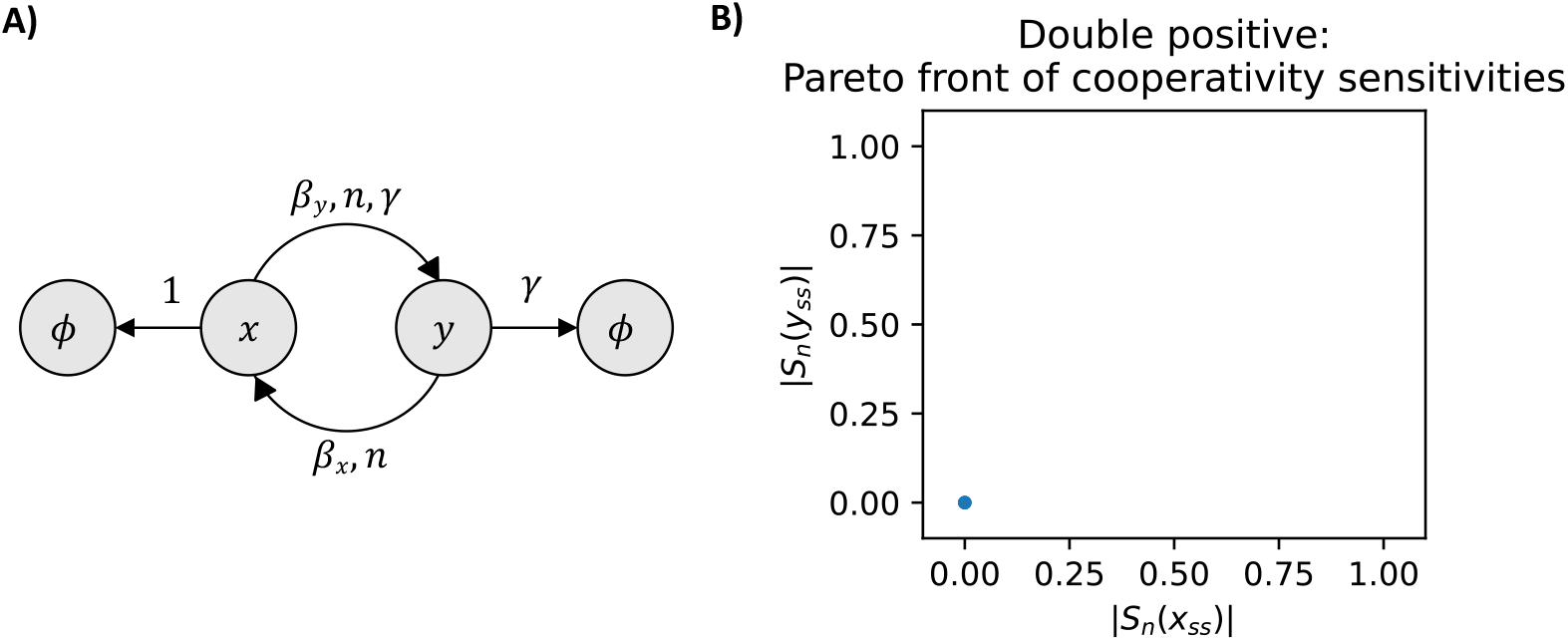
The diagram of the positive-positive feedback system and the Pareto front of a representative pair of sensitivity functions. **Panel A)** shows the diagram of the biological system with two positive feedbacks. The pointed arrows between species *x* and *y* represent positive feedback, where *x* enhances the production of *y* and vice versa. We assume that species *x* and *y* also undergo degradation and dilution within the cell, as indicated by the arrows pointed to the empty set symbol. The ratio of the rates of degradation and dilution is captured by the unitless parameter *γ*. After non-dimensionalization, the species *x* is degraded at a rate of 1, *β*_*x*_ is the rate of production of species *x* and *β*_*y*_*γ* is the rate of production of species *y*. Here, *n* represents the cooperativity of both species. **Panel B)** illustrates the Pareto front corresponding to the simultaneous minimisation of the absolute values of the sensitivities of species *x* and *y* with respect to the cooperativity *n*. The Pareto front for the positive-positive feedback system is the single point in blue. Solving the MOO problem shows that there is no trade-off between this pair of sensitivities. This result holds for all pairs of sensitivity functions, as plotted in Figure S3. The corresponding parameters are shown in Figure S4.

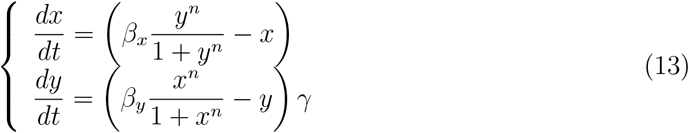

In this model, *β*_*x*_ and *β*_*y*_ are the ratios of the production rates to the degradation rates per unit of dissociation constants for species *x* and *y*, respectively. The constant *γ* is the ratio of the degradation rate of species *x* divided by the degradation rate of *y*. The constant *n* represents the cooperativity of the two species and we assume it to be the same for simplicity. We note that *β*_*x*_, *β*_*y*_, *γ*, and *n* are unitless and we refer to *β*_*x*_ and *β*_*y*_ as the strengths of the two positive feedbacks.

The steady states for this system, (*x*_*ss*_, *y*_*ss*_), occur when:

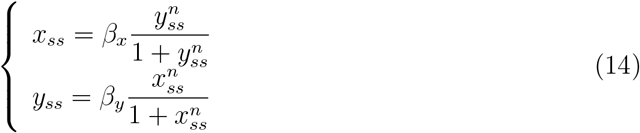

We note that the ratio of the degradation rates, *γ*, is not present in the steady-state equations. Plotting the nullclines in the parameter ranges that we consider biologically relevant reveals a single non-zero steady-state solution that is stable. It is the only solution we consider for this analysis.

The sensitivities functions of each steady-state coordinate to parameters *β*_*x*_, *β*_*y*_ and *n* are given in Table 2. We note that due to symmetry, it must be that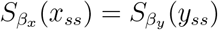. In Figure 3B), we illustrate a representative result of the problem of simultaneously minimising the absolute values of a pair of sensitivity functions. The Pareto front is comprised of a single point. This result holds across all pairs of sensitivity functions for combinations of both species (Supplementary Materials). Therefore, the double-positive feedback system is robust to changes in its biochemical parameters. This may be caused by the amplification created by the two positive feedbacks, mirroring the result of a single feedback system.

#### 2.3.4 The positive-negative feedback circuit is constrained by four trade-offs

The combination of positive and negative feedback is found in several biological processes such as the cell cycle. During the G2/M transition, positive feedback through cyclin-CDK complexes promotes rapid entry into mitosis, while negative feedback through checkpoint proteins such as p53 prevents progression if the DNA is damaged (*35*). The combination of positive and negative feedback is also believed to be responsible for robust oscillations found in natural systems. Synthetic circuits with these feedbacks were built in (*7, 36*).

In the positive-negative feedback system, one species *x* promotes the expression of another species *y*, which in turn inhibits the production of *x*. We propose a simple mathematical model to capture these interactions. If we use similar biochemical assumptions to those in the derivation of the model for the double-positive feedback system, the dynamics of the positive-negative feedback system can be described by Equation (15) and are illustrated in Figure 4A).

**Figure 4:**
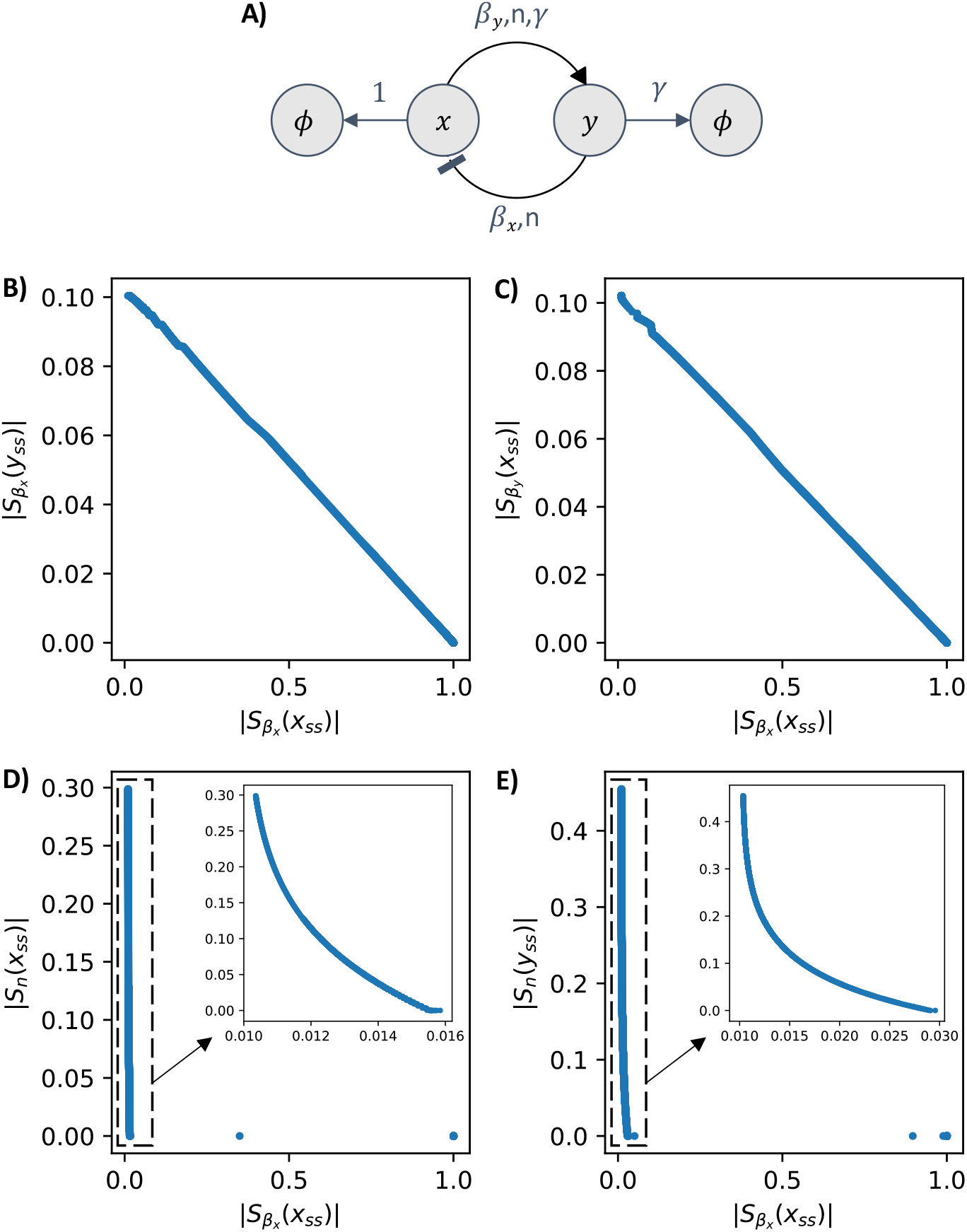
The diagram of the positive-negative feedback system and the Pareto fronts of four pairs of sensitivity functions. **Panel A)** shows the diagram of a biological system with positive feedback (pointed arrow) from species *x* to species *y*, and negative feedback (blunt arrow) from species *y* to species *x*. Both species undergo degradation. After non-dimensionalization, *x* is degraded at a rate of 1, *y* is degraded at a rate of *γ*, and production rates are *β*_*x*_ for *x* and *β*_*y*_*γ* for *y*. The binding cooperativity is *n*. **Panels B), C), D), and E)** display the Pareto fronts obtained from the simultaneous minimizations of the absolute sensitivity of species *x* to its production rate *β*_*x*_ against four other sensitivities. The four Pareto fronts, shown as blue lines, highlight trade-offs between these four pairs of sensitivities. The arrows in panels D) and E) point to a zoomed-in view of the Pareto fronts. Solving the multi-objective optimization problems reveals four distinct trade-offs in the positive-negative feedback system. Additional plots of Pareto-optimal sensitivities and their parameters are found in Figures S5 and S6.

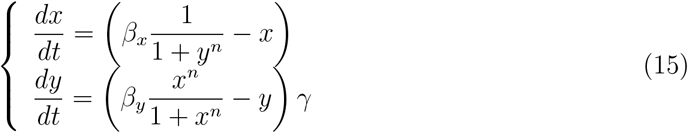

The steady states for this system, (*x*_*ss*_, *y*_*ss*_), occur when:

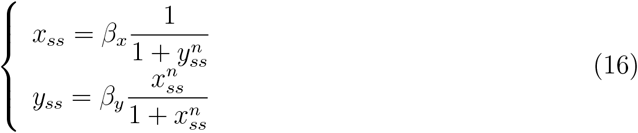

In this model, the nullclines intersect at a single steady-state solution, which is always non-zero unless *β*_*x*_ = 0. Thus, within our parameter ranges of interest, a single stable nonzero steady-state is found. We suspect that the model would require additional complexity to produce the oscillations observed in the literature.

For the single steady-state of the positive-negative system, we compute the sensitivity functions to variations in biochemical parameters and list them in Table 2. We note that due to symmetry, it must be that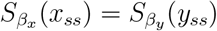. In Figure 4, we can observe trade-offs between the sensitivity function 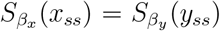 and all the other sensitivity functions: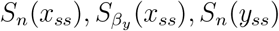, and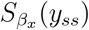. There are no trade-offs for other pairs of sensitivities. The simultaneous minimization of 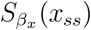 and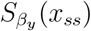, as well as 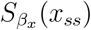 and *S*_*n*_(*x*_*ss*_) yields Pareto fronts at very high cooperativity values for *n*. Thus, to design a robust positive-negative feedback circuit, increasing the binding cooperativity as much as possible is the best design choice.

#### 2.3.5 The double-negative feedback loop (the toggle switch) is unconstrained by trade-offs

The toggle switch is a classic synthetic circuit in which two species mutually inhibit one another while individually undergoing degradation (*25*). The toggle switch is a “winner takes all” system, where one species ends up depleted and the other plateaus to a higher steady-state concentration. The toggle switch, illustrated in Figure 5, can be described by the model in Equation (17). This is a gene-interaction level model obtained from (*37*).

**Figure 5:**
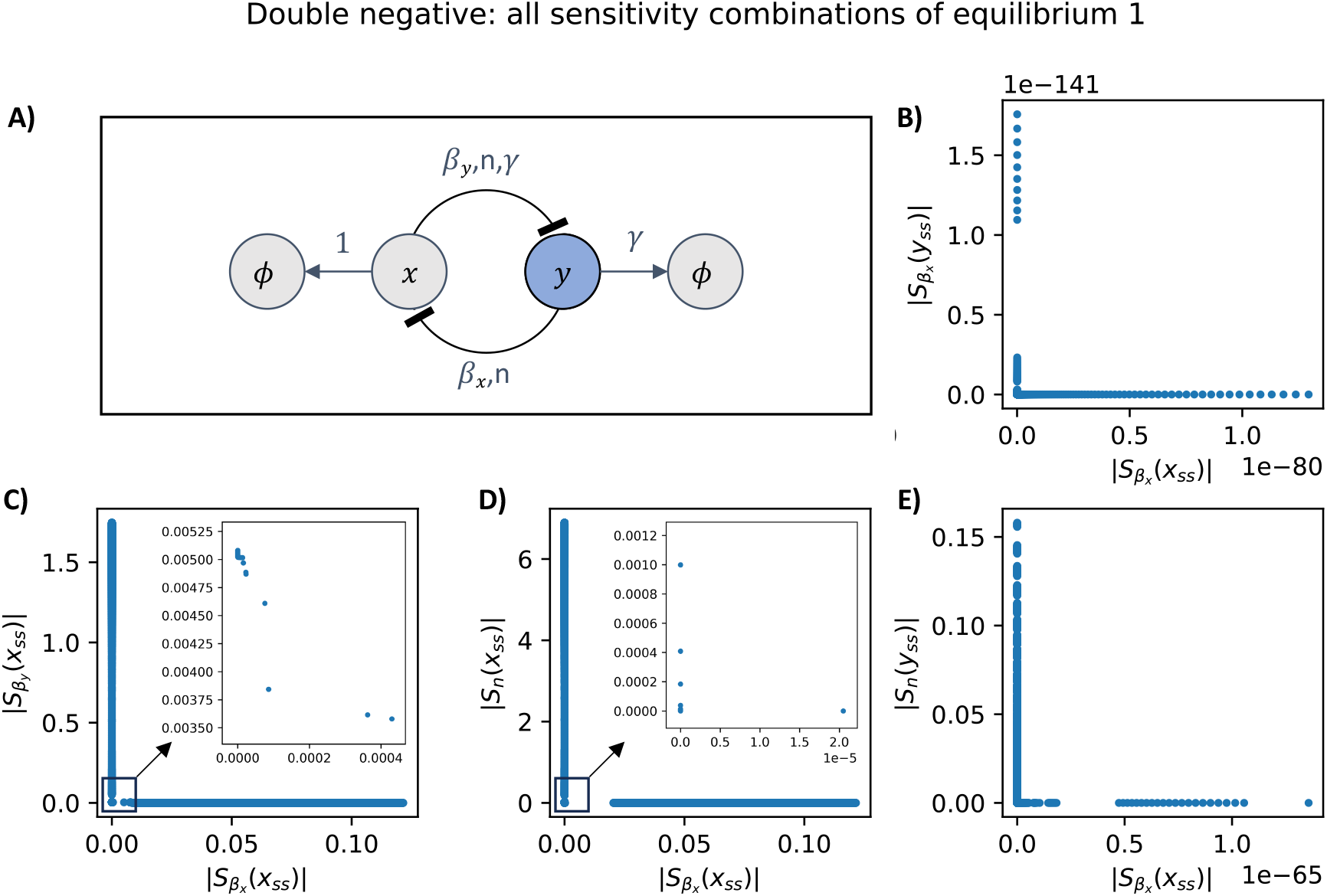
The diagram of the toggle switch and four of its Pareto fronts. **Panel A)** shows the diagram of two biochemical species *x* and *y* that mutually inhibit each other. The inhibitions are illustrated by blunt arrows. Both species *x* and *y* are subject to degradation, as indicated by arrows pointing to the empty set symbol. The reaction rates are listed above each arrow. **Panels B)-E)** show the Pareto fronts obtained by simultaneous minimisation of four sensitivity function pairs. In panels C) and D), we include the zoomed in version of the same figure. All four sensitivity function pairs are simultaneously minimised close to zero. Thus, no trade-offs are present in the model of the toggle switch. Plots for all other sensitivities combinations and their corresponding parameters are included in Figures S7-S10.

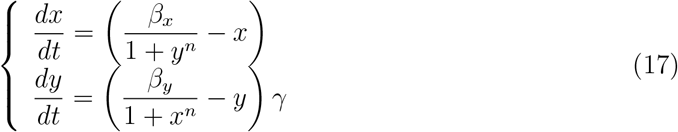

The steady-states for this system, (*x*_*ss*_, *y*_*ss*_), occur when:

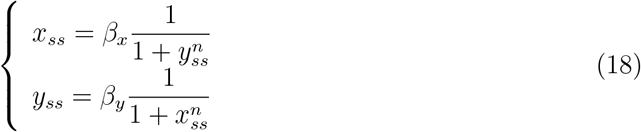

It is well-known that these two nullclines can intersect at up to three steady-state solutions. When three solutions are present, two are stable, and one is unstable. However, as the parameters change, the system undergoes a bifurcation where a stable and an unstable solution collide and disappear, leaving only a single stable solution. When only one stable steady-state exists, we compute its sensitivity function and later use it to supplement the two sensitivity functions that arise when two stable steady-states appear after the bifurcation.

The sensitivity functions associated with the stable steady-states of the toggle are given in Table 3. We note that due to symmetry, it must be that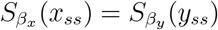. In Figure 5, we illustrate representative results of the problem of simultaneously minimising the absolute values of a pair of sensitivity functions. We direct the reader to the Supplementary Materials for the all the results of simultaneously minimising pairs of sensitivity functions. All of these MOO problems are solved to find Pareto fronts close to value zero. Thus, the toggle circuit is free of trade-offs and robust to variations in its biochemical parameters.

### 2.4 Promoter leakiness reduces trade-offs across all five circuits

Previously, we considered ideal scenarios where gene regulation is unaffected by promoter leakiness. However, in the presence of promoter leakiness, genes can still be expressed. Interestingly, our research reveals that promoter leakiness can enhance circuit robustness. In this section, we consider the case of promoter leakiness for the negative autoregulation circuit. We assume that the promoter’s leakiness rate can range between 0 and 20% of the maximum expression of the promoter (*38, 39*). This assumption should represent low and moderately leaky promoters. By denoting the promoter leakiness as the parameter *l*, the model of the negative autoregulation circuit is:

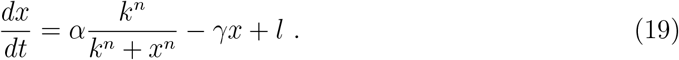

Using the change of variables in Table 1, we arrive at the non-dimensionalized model

**Table 1:**
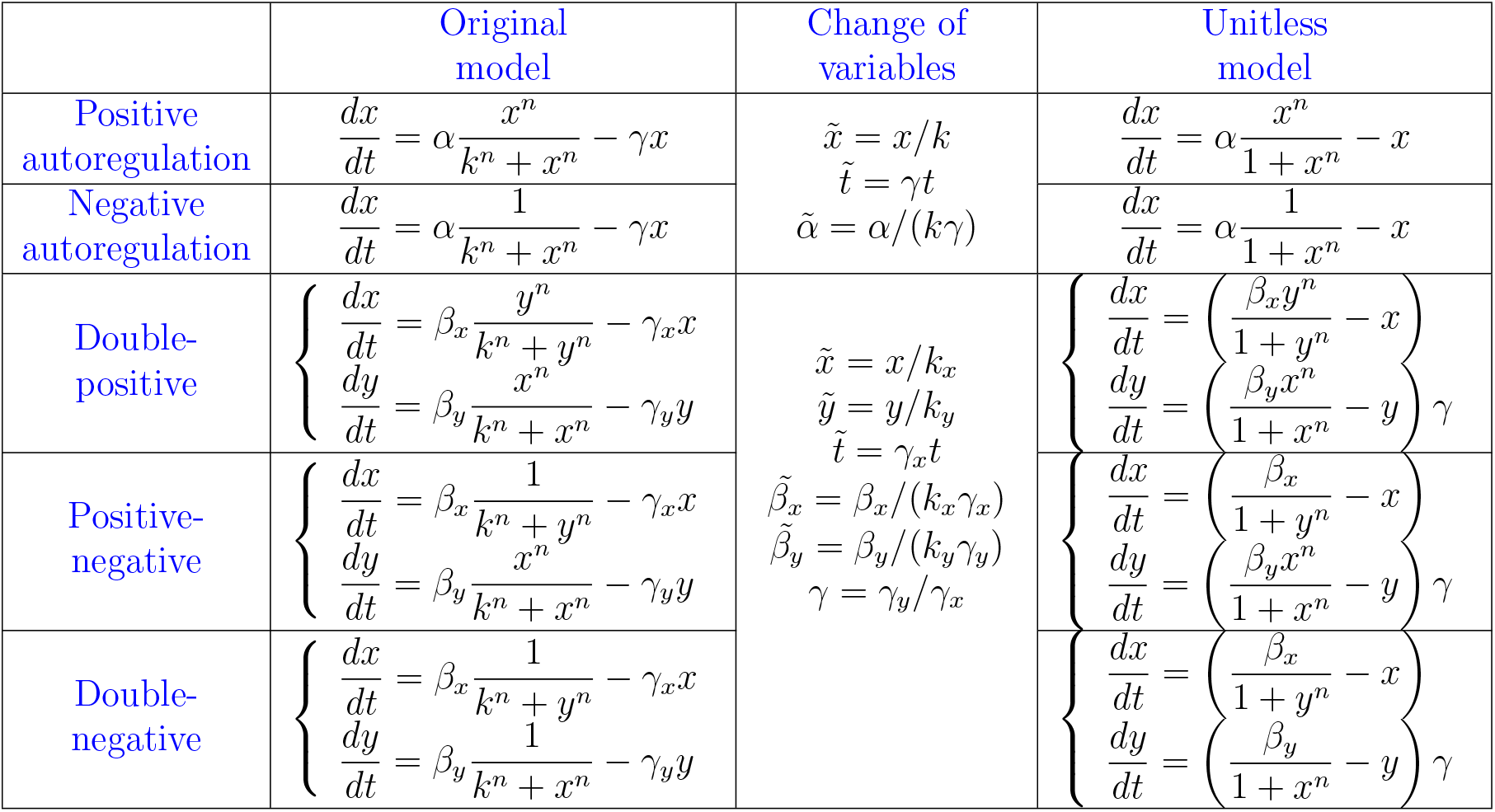
Mathematical models before and after non-dimensionalization for each of the biological systems studied in the paper. Tildes denote the unitless analogue of their original physical quantity. For simplicity, we use the variables without tildes in the unitless models.

|  | Original model | Change of variables | Unitless model |
| --- | --- | --- | --- |
| Positive autoregulation | $\frac{dx}{dt} = \alpha \frac{x^n}{k^n + x^n} - \gamma x$ | $\tilde{x} = x/k$<br>$\tilde{t} = \gamma t$<br>$\tilde{\alpha} = \alpha/(k\gamma)$ | $\frac{dx}{dt} = \alpha \frac{x^n}{1 + x^n} - x$ |
| Negative autoregulation | $\frac{dx}{dt} = \alpha \frac{1}{k^n + x^n} - \gamma x$ | | $\frac{dx}{dt} = \alpha \frac{1}{1 + x^n} - x$ |
| Double-positive | $\begin{cases} \frac{dx}{dt} = \beta_x \frac{y^n}{k^n + y^n} - \gamma_x x \\ \frac{dy}{dt} = \beta_y \frac{x^n}{k^n + x^n} - \gamma_y y \end{cases}$ | $\tilde{x} = x/k_x$<br>$\tilde{y} = y/k_y$<br>$\tilde{t} = \gamma_x t$<br>$\tilde{\beta}_x = \beta_x/(k_x \gamma_x)$<br>$\tilde{\beta}_y = \beta_y/(k_y \gamma_y)$<br>$\gamma = \gamma_y/\gamma_x$ | $\begin{cases} \frac{dx}{dt} = \left( \frac{\beta_x y^n}{1 + y^n} - x \right) \\ \frac{dy}{dt} = \left( \frac{\beta_y x^n}{1 + x^n} - y \right) \end{cases} \gamma$ |
| Positive-negative | $\begin{cases} \frac{dx}{dt} = \beta_x \frac{1}{k^n + y^n} - \gamma_x x \\ \frac{dy}{dt} = \beta_y \frac{x^n}{k^n + x^n} - \gamma_y y \end{cases}$ | | $\begin{cases} \frac{dx}{dt} = \left( \frac{\beta_x}{1 + y^n} - x \right) \\ \frac{dy}{dt} = \left( \frac{\beta_y x^n}{1 + x^n} - y \right) \end{cases} \gamma$ |
| Double-negative | $\begin{cases} \frac{dx}{dt} = \beta_x \frac{1}{k^n + y^n} - \gamma_x x \\ \frac{dy}{dt} = \beta_y \frac{1}{k^n + x^n} - \gamma_y y \end{cases}$ | | $\begin{cases} \frac{dx}{dt} = \left( \frac{\beta_x}{1 + y^n} - x \right) \\ \frac{dy}{dt} = \left( \frac{\beta_y}{1 + x^n} - y \right) \end{cases} \gamma$ |

**Table 2:** The sensitivity functions of the positive-positive feedback and the positive-negative feedback example systems.

|  |  |
| --- | --- |
| **Positive-Positive Feedback** | $S_{\beta_x}(x) = \frac{\beta_x \beta_y}{n^2 x (-y + \beta_y) + \beta_x (n^2 y - (-1 + n^2) \beta_y)}$ |
| | $S_{\beta_y}(x) = \frac{n \beta_y}{n^2 y + \left(1 - n^2 + \frac{x}{-x + \beta_x}\right) \beta_y}$ |
| | $S_n(x) = \frac{n (-ny \log[x] + (n \log[x] + \log[y]) \beta_y)}{n^2 y + \left(1 - n^2 + \frac{x}{-x + \beta_x}\right) \beta_y}$ |
| | $S_{\beta_x}(y) = \frac{n \beta_x}{n^2 x + \beta_x \left(1 - n^2 + \frac{y}{-y + \beta_y}\right)}$ |
| | $S_{\beta_y}(y) = \frac{\beta_x \beta_y}{n^2 x (-y + \beta_y) + \beta_x (n^2 y - (-1 + n^2) \beta_y)}$ |
| | $S_n(y) = \frac{n (-nx \log[y] + (\log[x] + n \log[y]) \beta_x)}{n^2 x + \beta_x \left(1 - n^2 + \frac{y}{-y + \beta_y}\right)}$ |
| **Positive-Negative Feedback** | $S_{\beta_x}(x) = \frac{\beta_x \beta_y}{n^2 (\beta_x - x) (\beta_y - y) + \beta_x \beta_y}$ |
| | $S_{\beta_y}(x) = -\frac{n \beta_y (\beta_x - x)}{n^2 (\beta_x - x) (\beta_y - y) + \beta_x \beta_y}$ |
| | $S_n(x) = -\frac{n (n \ln(x) x + \ln(y) \beta_y) (\beta_x - x)}{n^2 (\beta_x - x) (\beta_y - y) + \beta_x \beta_y}$ |
| | $S_{\beta_x}(y) = \frac{n \beta_x (\beta_y - y)}{n^2 (\beta_x - x) (\beta_y - y) + \beta_x \beta_y}$ |
| | $S_{\beta_y}(y) = \frac{\beta_x \beta_y}{n^2 (\beta_x - x) (\beta_y - y) + \beta_x \beta_y}$ |
| | $S_n(y) = \frac{n (\ln(x) \beta_x - n \ln(y) (\beta_x - x)) (\beta_y - y)}{n^2 (\beta_x - x) (\beta_y - y) + \beta_x \beta_y}$ |

**Table 3:** The sensitivity functions of the toggle switch.

|  |  |
| --- | --- |
| **Negative-Negative Feedback** | $S_{\beta_x}(x) = \frac{\beta_x \beta_y}{-n^2 (x - \beta_x) (y - \beta_y) + \beta_x \beta_y}$ |
| | $S_{\beta_y}(x) = \frac{n (x - \beta_x) \beta_y}{n^2 x (-y + \beta_y) + \beta_x (n^2 y - (-1 + n^2) \beta_y)}$ |
| | $S_n(x) = -\frac{n (x - \beta_x) (ny \log[x] + (-n \log[x] + \log[y]) \beta_y)}{-n^2 y \beta_x + n^2 x (y - \beta_y) + (-1 + n^2) \beta_x \beta_y}$ |
| | $S_{\beta_x}(y) = \frac{n \beta_x (y - \beta_y)}{n^2 x (-y + \beta_y) + \beta_x (n^2 y - (-1 + n^2) \beta_y)}$ |
| | $S_{\beta_y}(y) = \frac{\beta_x \beta_y}{-n^2 (x - \beta_x) (y - \beta_y) + \beta_x \beta_y}$ |
| | $S_n(y) = -\frac{n (nx \log[y] + (\log[x] - n \log[y]) \beta_x) (y - \beta_y)}{-n^2 y \beta_x + n^2 x (y - \beta_y) + (-1 + n^2) \beta_x \beta_y}$ |

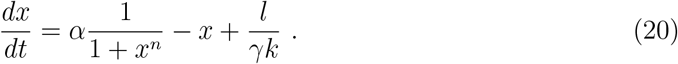

The quantity 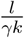 is a unitless value and so we can denote it as the non-dimensionalised leakiness parameter *L*. The equation reduces to:

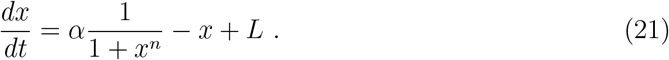

As outlined in the Methods section, the parameters *α, k* and *γ* have ranges of interest of [0.02, 0.1] nM min^−1^, [0.01, 10] nM and [0.01, 0.24] min^−1^, respectively. We have assumed that the leakiness rate *l* can range between 0 and 0.2*α* (*38*). Our range for the parameter *L* is therefore [0.002, 200].

In Figure 2, we observed that negative autoregulation was constrained by trade-offs. Figure 6 highlights the effects of introducing promoter leakiness. The original result of *L* = 0 is shown in blue. At *L* = 0.1, with the same sampling of parameter pair (*α, n*), the area spanned by the sensitivity pairs (*S*_*α*_, *S*_*n*_) decreases slightly, and the Pareto front collapses drastically at the orange points. With each successive increase in promoter leakiness L, the parameter region of the pair (*S*_*α*_, *S*_*n*_) shrinks further, and the corresponding Pareto fronts continue to move toward the origin, eventually eliminating trade-offs entirely.

**Figure 6:**
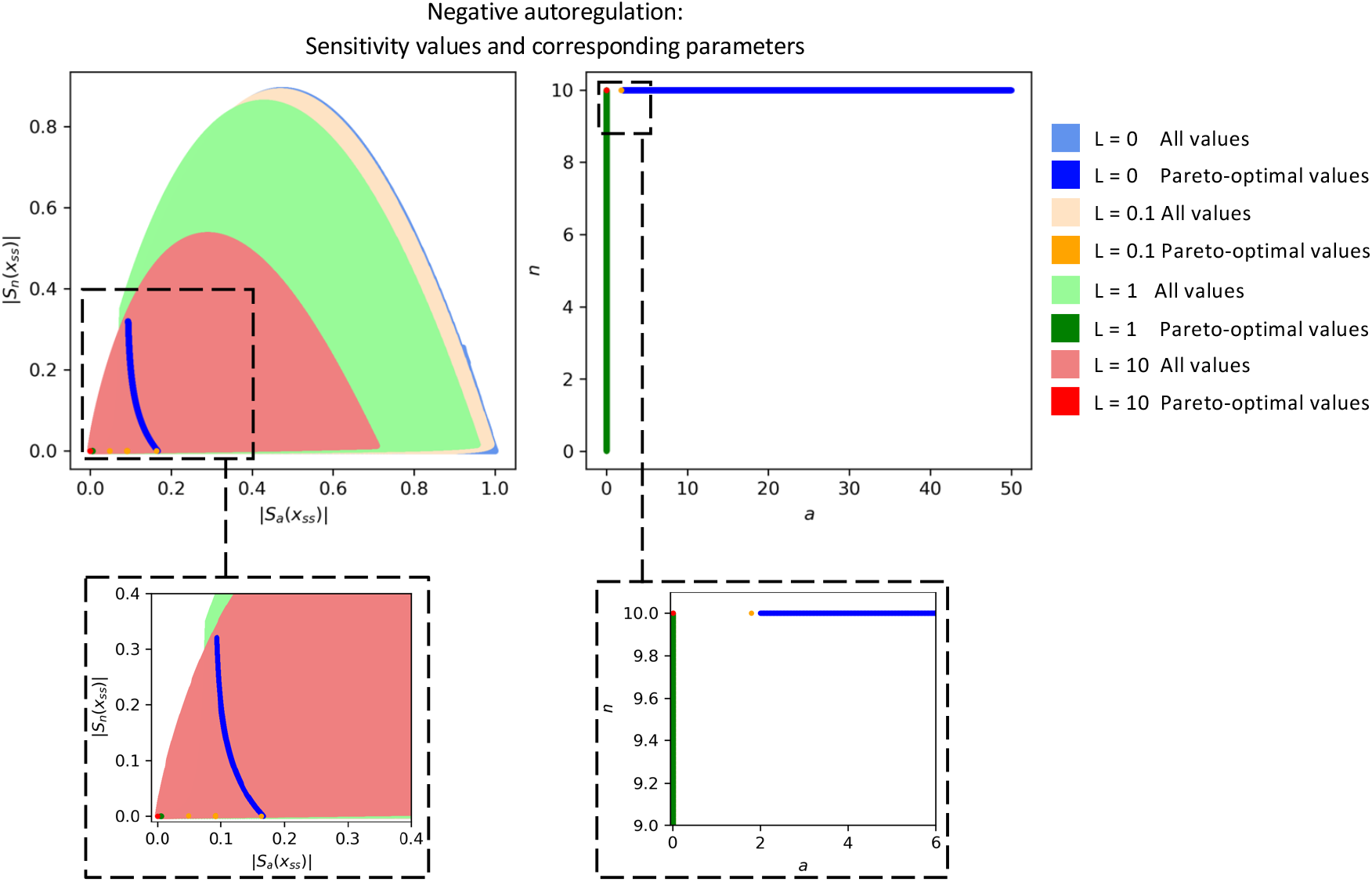
Increasing promoter leakiness enhances the robustness of the negative autoregulation circuit. We analyze four levels of leakiness: L=0 (blue), L=0.1 (orange), L=1 (green), and L=10 (red). In the left panel, sensitivity pairs (*S*_*α*_, *S*_*n*_) are plotted in lighter shades of their respective colors, forming contours that shrink as L increases. This shrinking parameter set illustrates how increased leakiness reduces sensitivity. Correspondingly, the Pareto fronts for each leakiness level also collapse toward the origin, with the smallest sensitivity values observed at the largest leakiness level (L=10). In the right panel, we show the parameter values corresponding to each Pareto front. Trade-off appears to occur when cooperativity is maximal whereas its absence seems to occur when the feedback strength is minimal.

This behavior can be explained as follows: as the leakiness constant L increases, the steady-state concentration of x rises. However, the rates *dx/dα* and *dx/dn* remain unaffected by the value of L. Consequently, the sensitivity functions *S*_*α*_(*x*_*ss*_) and *S*_*n*_(*x*_*ss*_) decrease. Thus, increasing leakiness actually renders negative autoregulation more robust.

The same pattern holds true for the other four circuits examined. In each case, increasing promoter leakiness collapses any existing trade-offs between pairs of sensitivity functions into a single point. For further details on the effects of promoter leakiness, we refer the reader to the supplementary information, where additional plots are provided.

Across all five synthetic circuits, we observed an absence of trade-offs between pairs of relative sensitivities when promoter leakiness is increased. Importantly, this result is not caused by the extensive optimization of the reaction rates corresponding to promoter strength, cooperativity, or production or degradation rates. By contrast, this result indicates that the combined regulatory mechanisms of feedback regulation and promoter leakiness inherently provide robustness and the decoupling of the sensitivities to variations in individual parameters. This result suggests that the combined regulatory mechanisms of feedback regulation and promoter leakiness provide flexibility to these circuits’ designs without the need for extensive optimization or fine-tuning.

We refer the reader to supplementary figures S11-S14 for plots illustrating the effect of promoter leakiness on the positive autoregulation circuit, the double-positive feedback circuit, the positive-negative feedback circuit, and the double-negative feedback circuit.

### 2.5 Adding a downstream gene to the positive autoregulation circuit does not introduce new trade-offs

We aim to explore the question of whether circuits that are unconstrained by trade-offs remain so if they are connected to downstream processes. As a case study, we connect the positive autoregulation circuit to a downstream gene and evaluate the impact on the trade-offs of the autoregulated gene’s sensitivity functions. In this analysis, we assume that the downstream gene does not use up the species in the positive feedback circuit. The result is that the positive autoregulation circuit remains unconstrained by trade-offs after the addition of the downstream gene.

To demonstrate this, we compare the trade-offs for a positively autoregulated gene x with production rate (*α*) and cooperativity (*n*) to a scenario where x produces an external gene y without incurring any cost to the x population. This comparison is depicted in Figure 7B, where the circuitry represented by solid lines corresponds to the autoregulation of x, and the circuitry represented by dashed lines illustrates the downstream components. The former represents the previously established positive autoregulation circuit, while the latter can be mathematically described as follows:

**Figure 7:**
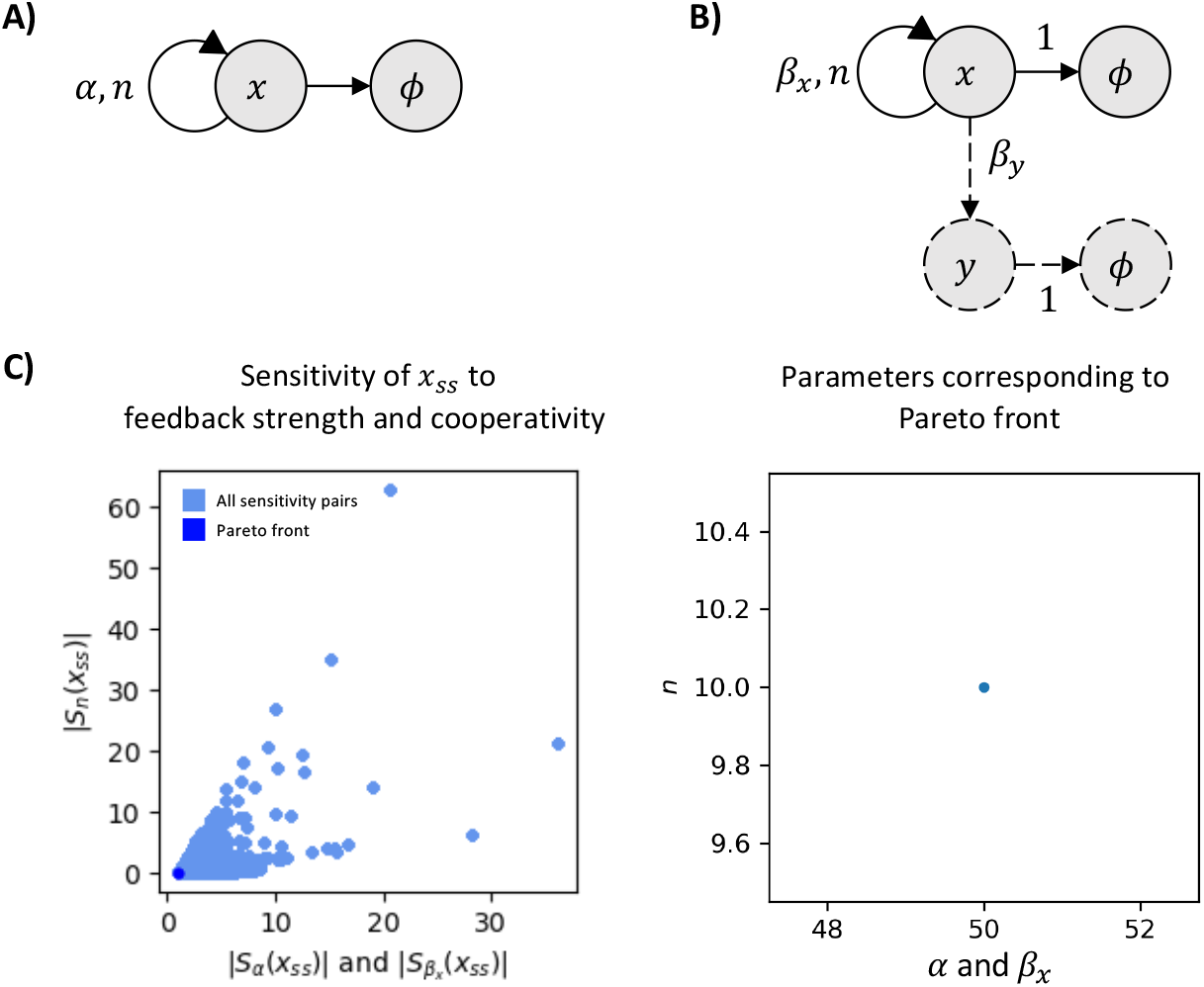
Adding a downstream gene to the positive feedback circuit does not introduce any new trade-offs. **Panel A)** Diagram of a single positive autoregulating species. **Panel B)** The same positive autoregulation circuit (drawn in solid lines) regulates a downstream gene (drawn in dashed lines). **Panel C)** (Left) Plot of all sensitivity values (light blue) and their Pareto front (dark blue) of sensitivity of x steady state to its feedback strength (*α* for circuit in panel A and *β*_*x*_ for circuit in panel B) and to its cooperativity constant (*n*). A single Pareto point at (1, 0) is observed, indicating no tradeoff. (Right) The corresponding parameter for this Pareto point is at maximal feedback strength and maximal cooperativity: *α, β*_*x*_ = 50 and *n* = 10. The circuits in panels A) and B) share the same sensitivity functions.

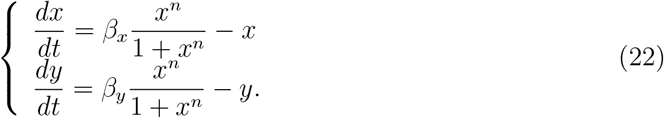

Here, the rate *β*_*x*_ represents the feedback strength of x (identical to *α*) and *β*_*y*_ represents the production rate of y driven by x. Both circuits have the same sensitivity values with respect to their respective feedback strength and cooperativity. The sensitivity space is therefore identical and so are their Pareto fronts at 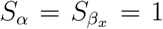 and *S*_*n*_ = 0. Since the inclusion of downstream gene y does not use up the species in the positive feedback circuit, the sensitivities and trade-offs of the combined circuit remain unaffected.

### 2.6 Positive autoregulation can introduce new trade-offs in down-stream gene circuits, but it can also enhance their robustness

As a case study on the impact of incorporating feedback into a circuit, we examine a two-species system in which gene X regulates gene Y, and explore the effect of changing the expression of X from constitutive expression to positive autoregulation. We illustrate these circuits in Figure 8, panels (A) and (D). The dynamics of the modified circuit are governed by the equations

**Figure 8:**
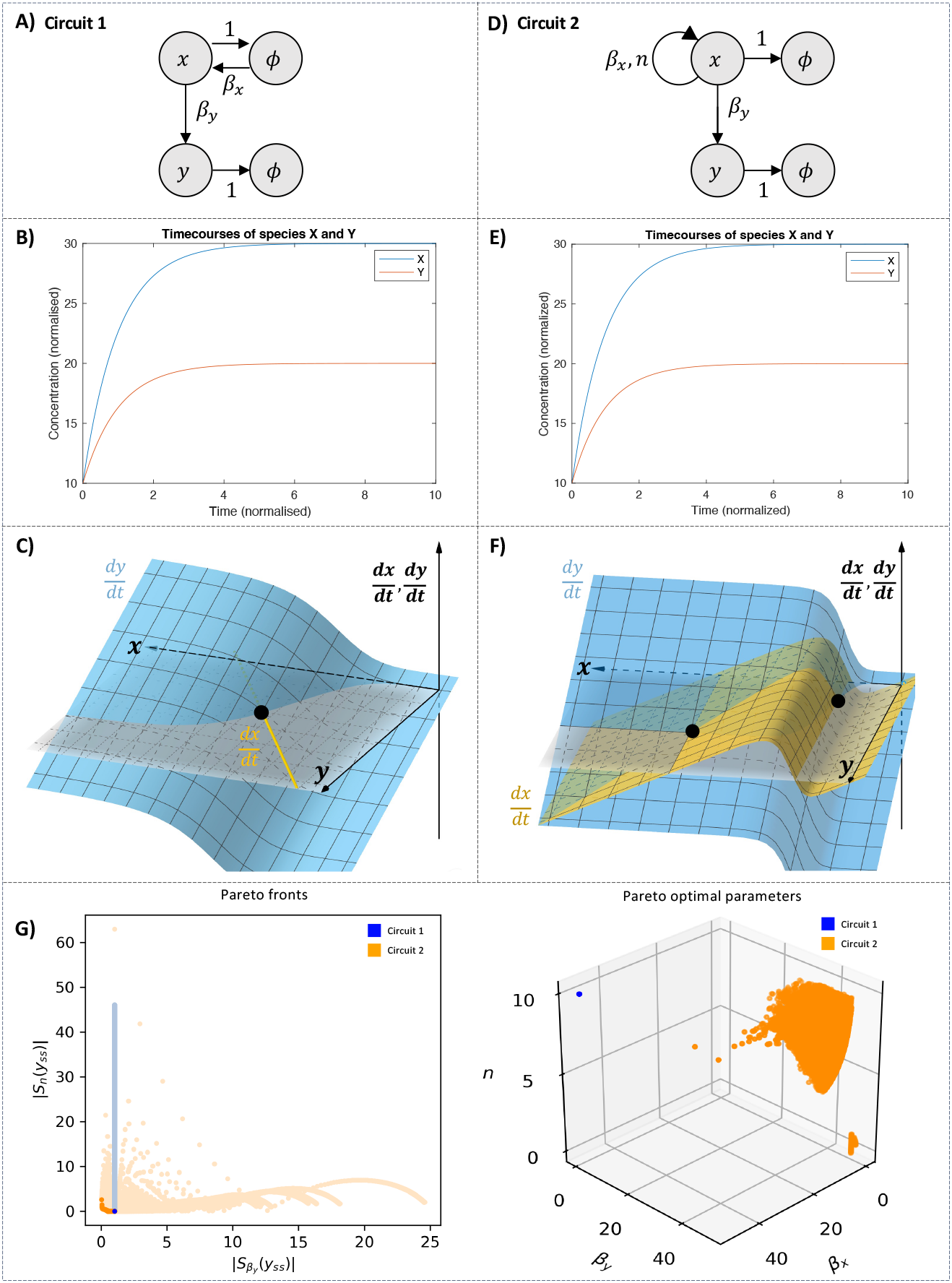
Introducing positive feedback upstream can create trade-offs for genes expressed downstream. **Panel A)** depicts a constitutively expressed gene X regulating a gene Y. The rate of X expression is denoted by *β*_*x*_ and the rate of Y expression is denoted by *β*_*y*_. In **Panel D)**, the constitutive expression of X is replaced with positive autoregulation of feedback strength *β*_*x*_ and cooperativity *n*. **Panels B) and E)** plot the timecourses of species x and y for identical conditions: *x*(0) = 10, *y*(0) = 10, *β*_*x*_ = 30, *β*_*y*_ = 20, and *n* = 10. Both circuits settle into a stable steady-state. **Panels C) and F)** show that Circuit 1 only has one steady state (C) while Circuit 2 can have up to two (F). **Panel G)** contains two plots. The left hand plot shows the span of sensitivity function pairs 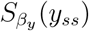 and *S*_*n*_(*y*_*ss*_) and their Pareto fronts. Circuit 1 is shown in blue and Circuit 2 in orange. Introducing positive autoregulation for species X creates a new trade-off (dark orange curve). The corresponding parameters for the Pareto front are shown in the right hand plot.

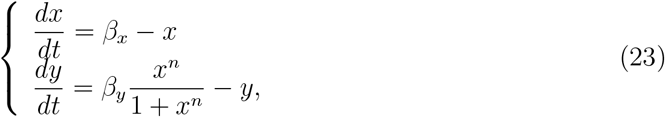

while the dynamics of the latter are the same as in Equation (22).

Without positive autoregulation, the sensitivities of the steady-state concentration of species y to its production rate (*β*_*y*_) and cooperativity constant (*n*) are unconstrained by a trade-off. The system exhibits a single optimal choice of parameters at which it is maximally robust. This is illustrated in blue in Figure 8 (G), where the light blue points in the left panel represent all sensitivities accessible to the system, and the dark blue point represents the Pareto front. When positive autoregulation is incorporated into dynamics of gene *x*, trade-offs emerge for the sensitivities of the steady-state species y with respect to *β*_*y*_ and *n*. This is shown in orange, where the light orange points in the left panel represent the new sensitivities spanned by sampling the same parameter space, and the dark orange curve represents the resulting Pareto front. This new Pareto front demonstrates that a trade-off now exists between sensitivity to the production rate and cooperativity. The system with positive autoregulation can access greater robustness to production rate than in the previous case, but at the expense of increased sensitivity to cooperativity. Our findings suggest that positive autoregulation can enhance the robustness of downstream genes to one parameter variation at the expense of another, by introducing trade-offs that can access regions closer to the origin.

## 3 CONCLUSION

In this paper, we introduced a general computational framework to analyse how robust biological systems with feedback regulation are to variations in their biochemical parameters and we revealed the trade-offs in robustness that are present for these systems. Our computational framework used multi-objective optimization to simultaneously minimize the sensitivities of these biological systems to variations in pairs of biochemical parameters. The results of the multi-objective optimization problems revealed whether trade-offs between the robustness to variations in pairs of parameters were present.

To illustrate this computational framework, we considered five examples of biological systems with feedback and analysed their sensitivity to variations in biochemical parameters. In single-species feedback, a trade-off between the sensitivity to feedback strength and to cooperativity was found for negative autoregulation but was absent for positive autoregulation. Indeed, we found that in negative autoregulation, increasing the robustness to variations in cooperativity was only possible at a loss of robustness in feedback strength, and vice versa. In dual-species feedback, we found several trade-offs between the strengths of feedback and the cooperativity of biomolecules. In double-positive feedback and the toggle switch, no trade-offs arose, similar to positive autoregulation. However, positive-negative feedback was constrained by trade-offs, requiring that the system design carefully balances robustness to feedback strength and cooperativity.

Although each of the five circuits had distinct functional roles, common features shared among them were observed. The net feedback sign of a circuit appeared to be related to the presence or absence of trade-offs. The net-positive feedback architectures (e.g., positive autoregulation, double-positive feedback, and double-negative feedback) were interrelated by their lack of trade-offs in sensitivity functions, while the net-negative architectures (e.g., negative autoregulation and positive-negative feedback) were interrelated by their trade-off constraints. This suggested that further research on more general net-positive feedback architectures might be fruitful for finding robust designs. Additionally, promoter leakiness was found to universally reduce trade-offs if they were present or to maintain the absence of trade-offs if none existed. The combination of promoter leakiness and feedback regulation demonstrated exceptional robustness and decoupled the sensitivities to variations in individual reaction rate parameters.

Overall, our work contributes to understanding trade-offs in the robustness of biological systems with feedback regulation. While it is well-known that engineered feedback is constrained by robustness trade-offs, as it cannot fully counteract all disturbances, similar insights for biological feedback systems remain underexplored. This paper helps bridge that gap by providing evidence of analogous trade-offs in biological contexts.

Future directions of this research include optimizing the sampling algorithm to efficiently solve multiple-objective optimization problems for parameter spaces in higher than three dimensions. We discuss the limitations of the grid-sampler and potential other algorithms in Section 4.4 of the Methods. Further research could replace the sensitivity functions evaluated at steady-state with the sensitivity functions for transient dynamics developed by Wu *et al*. (*40*) and assess the robustness of biological systems such as oscillators (*7, 41*). Lastly, further research could use this computational framework to research other examples of biological feedback systems and to develop theoretical arguments for why some examples of feedback introduce robustness trade-offs, while others do not.

Our current research contributes to advancing the understanding of biological feedback in natural systems and to improving system design for synthetic biological circuits. Ultimately, we aim for our work to facilitate future research aimed at unraveling the complexities of feedback in biological systems and at enhancing the reliability of engineered biological applications.

## 4 METHODS

### 4.1 Non-dimensionalization of the mathematical models

Nondimensionalising biochemical kinetics involves defining the relationships between physical quantities with units and their unitless analogues, and performing a change of variables of the original equations. Table 1 lists the model pairs for each feedback mechanism, and the associated changes of variables.

### 4.2 Parameter ranges for the models of the biological systems

In this work, we assume that the synthesis rates *α, β* ∈ [0.02, 0.1] nM min^−1^ according to values from (*42*), that dissociation constants *k* ∈ [0.01, 10] nM (*42, 43*), and that the degradation and dilution rates *γ* ∈ [0.01, 0.24] min^−1^, as follows from the typical doubling times of bacterial and yeast cells (*44, 45*).

From Table 1, unitless feedback strengths 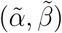 are defined as the ratio between the maximal production rate (*α, β*) and the product of dissociation constant (*k*) with degradation rate (*γ*). The range of values for the unitless feedback strengths is therefore 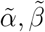 ∈ [1*/*120, 1000].

For practicality, we sample the range [0.01, 50].

Cooperative binding falls into four classes: binding that does not change the system dynamics (*n* = 0), binding that impedes further binding (0 *< n <* 1), binding that has no effect on subsequent binding (*n* = 1) and binding that promotes further binding (*n >* 1). At *n* = 0, the feedback disappears. Therefore, we consider *n >* 0. We set a lower bound of *n* = 0.01 for our numerical simulations. The upper bound was set to *n* = 10 to cover the single digit cooperativities that are common to biological systems (*46*).

### 4.3 Numerical methods

The equations for steady-states in Table 1 cannot typically be solved analytically to find their solutions *x*_*ss*_ and *y*_*ss*_. Instead, we employ scipy.optimize.fsolve from Python to solve these equations numerically. The parameter set we use was obtained by evenly grid-sampling the parameter ranges of *α, β* and *n*. Additionally, since we are only interested in the stable steady-states of our biological systems, we filtered out the unstable steady-states by imposing the condition that the real part of their eigenvalues evaluated at the solutions is less than zero. The resulting solutions and their respective parameter values were then used to evaluate the corresponding sensitivity functions via the analytically derived formulas. We refer the reader to the linked Mathematica files for the analytical derivations of the sensitivity functions.

In the MOO problems, the Pareto front was found using the Python package paretoset, based on the method proposed by (*47*).

### 4.4 Limitations of the sampling algorithm

The correct identification of the Pareto front is dependent on sufficiently sampling the two- or three-dimensional biochemical parameter space. The grid-search sampler we use is a computationally expensive method. For example, to find the Pareto fronts for the MOO problems in the positive-negative feedback system, we use 10^9^ samples for the three-dimensional grid. The development of computationally efficient algorithms for solving multi-objective optimization problems remains an active area of research. More efficient methods such as Multi-Objective Simulated Annealing (*48*) could be used in future work to reduce the number of samples required to find the Pareto fronts.

#### 4.5 Sensitivity functions

In this section, we include the analytical form of the sensitivity functions for all two-species examples in Tables 2 and 3.

For brevity, we refer the reader to the code supplied in the GitHub repository for all sensitivity functions associated with leakiness from Section 2.4, and additional circuits covered in Sections 2.5 and 2.6.

## Abbreviations

MOO: multi-objective optimization

## 5 Associated Content

### Data Availability Statement

The Supporting Information includes additional figures as listed below. The manuscript figures, the Supporting Information figures, and the code used for figure generation are available on GitHub at the following link: https://github.com/nguyenhntran/ACSPaperSep2024.

### Supporting Information

Pareto fronts of double positive feedback (Figure S1); Pareto-optimal parameters of double positive feedback (Figure S2); Pareto fronts of positive-negative feedback (Figure S3); Pareto-optimal parameters of positive-negative feedback (Figure S4); Pareto fronts of double negative feedback at first equilibrium (Figure S5); Pareto-optimal parameters of double negative feedback at first equilibrium (Figure S6); Pareto fronts of double negative feedback at second equilibrium (Figure S7); Pareto-optimal parameters of double negative feedback at second equilibrium (Figure S8); Effect of promoter leakiness on positive autoregulation (Figure S9); Effect of promoter leakiness on double positive feedback (Figure S10); Effect of promoter leakiness on positive-negative feedback (Figure S11); Effect of promoter leakiness on double-negative feedback (Figure S12) (PDF).

## 6 Author Information

### Corresponding Author

Dr. Ania-Ariadna Baetica, Department of Mechanical Engineering and Mechanics, Drexel University, Philadelphia, PA 19104, USA.

### Author Contributions

A.-A.B. conceptualized the project. N.H.N.T., A.N., and T.W.R. carried out the computational simulations. N.H.N.T. and A.-A.B. analyzed the results and wrote the manuscript.

### Notes

The authors declare no competing financial interest. The authors declare no funding sources.

## Notes

https://github.com/nguyenhntran/ACSPaperSep2024

### Competing Interest Statement

The authors have declared no competing interest.

### Summary of Updates

Figure 2 was revised. New Figures 6-8 have been added and their message was explained in Sections 2.4-2.6. The conclusion was updated with information from new Figures 6-8. Supplemental information was revised.

## References

1. Alon, U. An Introduction to Systems Biology: Design Principles of Biological Circuits; CRC press, 2019.

2. Del Vecchio, D., and Murray, R. M. Biomolecular Feedback Systems; Princeton University Press Princeton, NJ, 2015.

3. El-Samad, H., Goff, J., and Khammash, M. (2002) Calcium homeostasis and parturient hypocalcemia: an integral feedback perspective. Journal of Theoretical Biology 214, 17– 29.

4. Rosenfeld, N., Elowitz, M. B., and Alon, U. (2002) Negative autoregulation speeds the response times of transcription networks. Journal of Molecular Biology 323, 785–793.

5. Yi, T.-M., Huang, Y., Simon, M. I., and Doyle, J. (2000) Robust perfect adaptation in bacterial chemotaxis through integral feedback control. Proceedings of the National Academy of Sciences 97, 4649–4653.

6. Werner, J. (2010) System properties, feedback control and effector coordination of human temperature regulation. European Journal of Applied Physiology 109, 13–25.

7. Tsai, T. Y.-C., Choi, Y. S., Ma, W., Pomerening, J. R., Tang, C., and Ferrell Jr, J. E. (2008) Robust, tunable biological oscillations from interlinked positive and negative feedback loops. Science 321, 126–129.

8. Lahav, G. (2008) Oscillations by the p53-Mdm2 feedback loop. Cellular Oscillatory Mechanisms 28–38.

9. Baetica, A.-A., Westbrook, A., and El-Samad, H. (2019) Control theoretical concepts for synthetic and systems biology. Current Opinion in Systems Biology 14, 50–57.

10. El-Samad, H. (2021) Biological feedback control—Respect the loops. Cell Systems 12, 477–487.

11. Åström, K. J., and Murray, R. Feedback systems: an introduction for scientists and engineers; Princeton university press, 2021.

12. Doyle, J. C., Francis, B. A., and Tannenbaum, A. R. Feedback Control Theory; Courier Corporation, 2013.

13. Stein, G. (2003) Respect the unstable. IEEE Control Systems Magazine 23, 12–25.

14. Kitano, H. (2004) Biological robustness. Nature Reviews Genetics 5, 826–837.

15. Chandra, F. A., Buzi, G., and Doyle, J. C. (2011) Glycolytic oscillations and limits on robust efficiency. Science 333, 187–192.

16. El-Samad, H., Kurata, H., Doyle, J., Gross, C., and Khammash, M. (2005) Surviving heat shock: control strategies for robustness and performance. Proceedings of the National Academy of Sciences 102, 2736–2741.

17. Bhatnagar, R., and El-Samad, H. (2016) Tradeoffs in adapting biological systems. European Journal of Control 30, 68–75.

18. Olsman, N., Baetica, A.-A., Xiao, F., Leong, Y. P., Murray, R. M., and Doyle, J. C. (2019) Hard limits and performance tradeoffs in a class of antithetic integral feedback networks. Cell Systems 9, 49–63.

19. Baetica, A.-A., Leong, Y. P., and Murray, R. M. (2020) Guidelines for designing the antithetic feedback motif. Physical Biology 17.

20. Szekely, P., Sheftel, H., Mayo, A., and Alon, U. (2013) Evolutionary tradeoffs between economy and effectiveness in biological homeostasis systems. PLoS Computational Biology 9.

21. Kirschner, M. W., and Gerhart, J. C. The plausibility of life: Resolving Darwin’s dilemma; Yale University Press, 2005.

22. Shakiba, N., Jones, R. D., Weiss, R., and Del Vecchio, D. (2021) Context-aware synthetic biology by controller design: Engineering the mammalian cell. Cell systems 12, 561–592.

23. Yeung, E., Dy, A. J., Martin, K. B., Ng, A. H., Del Vecchio, D., Beck, J. L., Collins, J. J., and Murray, R. M. (2017) Biophysical constraints arising from compositional context in synthetic gene networks. Cell systems 5, 11–24.

24. Khammash, M. (2016) An engineering viewpoint on biological robustness. BMC Biology 14, 22.

25. Gardner, T. S., Cantor, C. R., and Collins, J. J. (2000) Construction of a genetic toggle switch in Escherichia coli. Nature 403, 339–342.

26. Shopera, T., Henson, W. R., Ng, A., Lee, Y. J., Ng, K., and Moon, T. S. (2015) Robust, tunable genetic memory from protein sequestration combined with positive feedback. Nucleic Acids Research 43, 9086–9094.

27. Stricker, J., Cookson, S., Bennett, M. R., Mather, W. H., Tsimring, L. S., and Hasty, J. (2008) A fast, robust and tunable synthetic gene oscillator. Nature 456, 516–519.

28. Rabitz, H., Kramer, M., and Dacol, D. (1983) Sensitivity analysis in chemical kinetics. Annual review of physical chemistry 34, 419–461.

29. Otero-Muras, I., and Banga, J. R. (2017) Automated design framework for synthetic biology exploiting pareto optimality. ACS Synthetic Biology 6, 1180–1193.

30. Hu, C. Y., and Murray, R. M. (2022) Layered feedback control overcomes performance trade-off in synthetic biomolecular networks. Nature communications 13, 5393.

31. Shoval, O., Sheftel, H., Shinar, G., Hart, Y., Ramote, O., Mayo, A., Dekel, E., Kavanagh, K., and Alon, U. (2012) Evolutionary trade-offs, Pareto optimality, and the geometry of phenotype space. Science 336, 1157–1160.

32. Jedlicka, P., Bird, A. D., and Cuntz, H. (2022) Pareto optimality, economy–effectiveness trade-offs and ion channel degeneracy: improving population modelling for single neurons. Open Biology 12.

33. Yamane, H., and Paul, W. E. (2012) Memory CD4+ T cells: fate determination, positive feedback and plasticity. Cellular and Molecular Life Sciences 69, 1577–1583.

34. Chang, D.-E., Leung, S., Atkinson, M. R., Reifler, A., Forger, D., and Ninfa, A. J. (2010) Building biological memory by linking positive feedback loops. Proceedings of the National Academy of Sciences 107, 175–180.

35. Hoffmann, I., and Karsenti, E. (1994) The role of cdc25 in checkpoints and feedback controls in the eukaryotic cell cycle. Journal of Cell Science 1994, 75–79.

36. Atkinson, M. R., Savageau, M. A., Myers, J. T., and Ninfa, A. J. (2003) Development of genetic circuitry exhibiting toggle switch or oscillatory behavior in Escherichia coli. Cell 113, 597–607.

37. Bois, J. S. justinbois/biocircuits: Version 0.1.0. 2021; 10.22002/D1.1615.

38. Ede, C., Chen, X., Lin, M.-Y., and Chen, Y. Y. (2016) Quantitative analyses of core promoters enable precise engineering of regulated gene expression in mammalian cells. ACS synthetic biology 5, 395–404.

39. Garcia, H. G., and Phillips, R. (2011) Quantitative dissection of the simple repression input–output function. Proceedings of the National Academy of Sciences 108, 12173– 12178.

40. Wu, W. H., Wang, F. S., and Chang, M. S. (2008) Dynamic sensitivity analysis of biological systems. BMC bioinformatics 9, 1–17.

41. Elowitz, M. B., and Leibler, S. (2000) A synthetic oscillatory network of transcriptional regulators. Nature 403, 335–338.

42. Kofahl, B., and Klipp, E. (2004) Modelling the dynamics of the yeast pheromone pathway. Yeast 21, 831–850.

43. Chen, K. C., Csikasz-Nagy, A., Gyorffy, B., Val, J., Novak, B., and Tyson, J. J. (2000) Kinetic analysis of a molecular model of the budding yeast cell cycle. Molecular biology of the cell 11, 369–391.

44. Helmstetter, C. E., and Cooper, S. (1968) DNA synthesis during the division cycle of rapidly growing Escherichia coli Br. Journal of molecular biology 31, 507–518.

45. Sherman, F. Methods in enzymology; Elsevier, 2002; Vol. 350; pp 3–41.

46. Bellelli, A. (2010) Hemoglobin and cooperativity: Experiments and theories. Current Protein and Peptide Science 11, 2–36.

47. Borzsony, S., Kossmann, D., and Stocker, K. The skyline operator. Proceedings 17th international conference on data engineering. 2001; pp 421–430.

48. Serafini, P. Simulated annealing for multi objective optimization problems. Multiple Criteria Decision Making: Proceedings of the Tenth International Conference: Expand and Enrich the Domains of Thinking and Application. 1994; pp 283–292.

