## Supplemental Materials for "Fundamental trade-offs in the robustness of biological systems with feedback regulation"

### 1 SUPPORTING INFORMATION

#### 1.1 Code Availability

The code for modeling all the examples of biological systems and their analysis is available on Github at `https://github.com/nguyenhntran/ACSPaperSep2024`.

#### 1.2 Supporting Figures

For all two species architectures, we have $S_{\beta_x}(x_{ss}) = S_{\beta_y}(y_{ss})$. There are a total of 10 unique sensitivity combinations out of a total 15 possible. We present them for each circuit in the following figures.

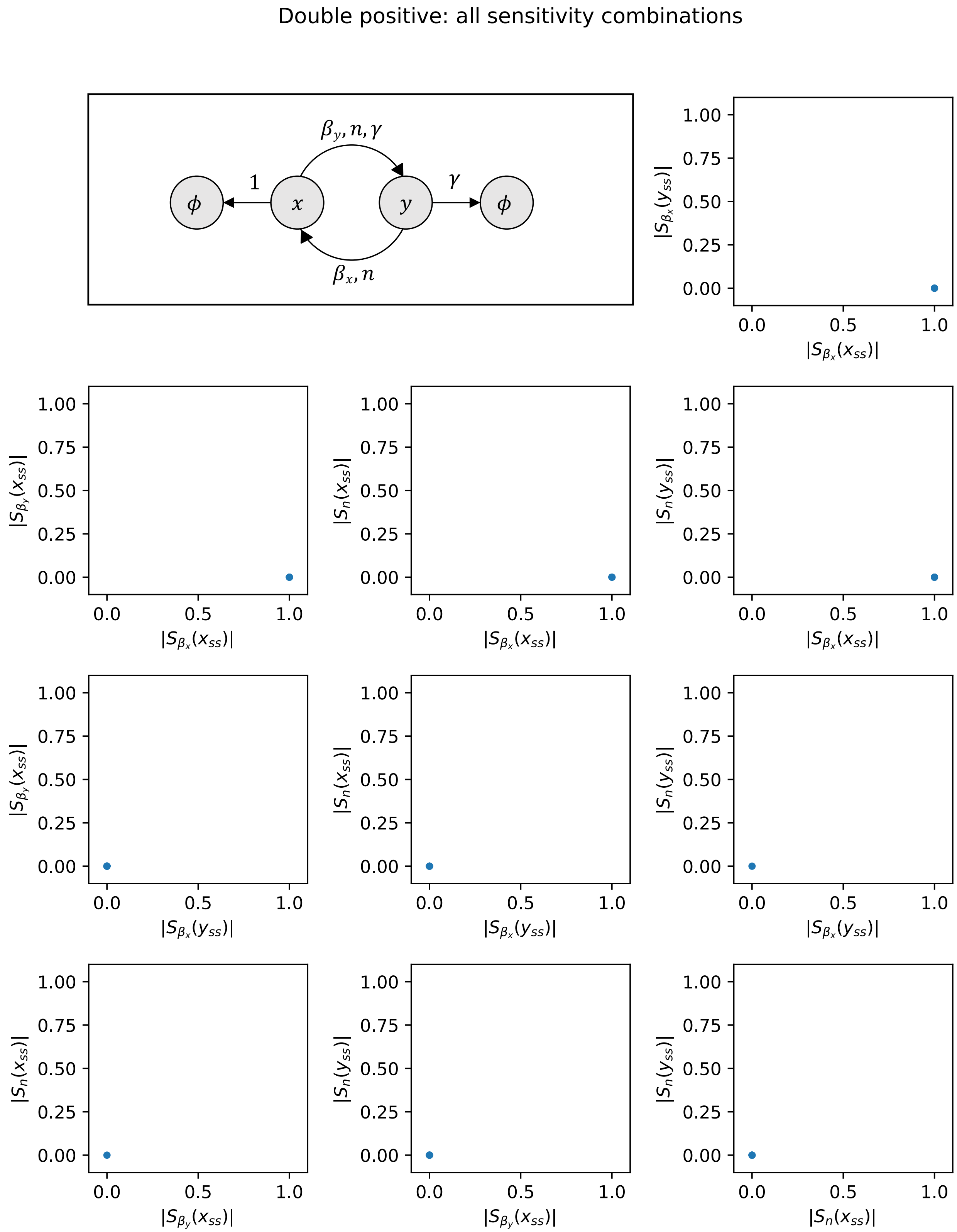


Figure S1: Pareto fronts obtained from the simultaneous minimization of all pairs of sensitivity combinations for the positive-positive feedback loop.

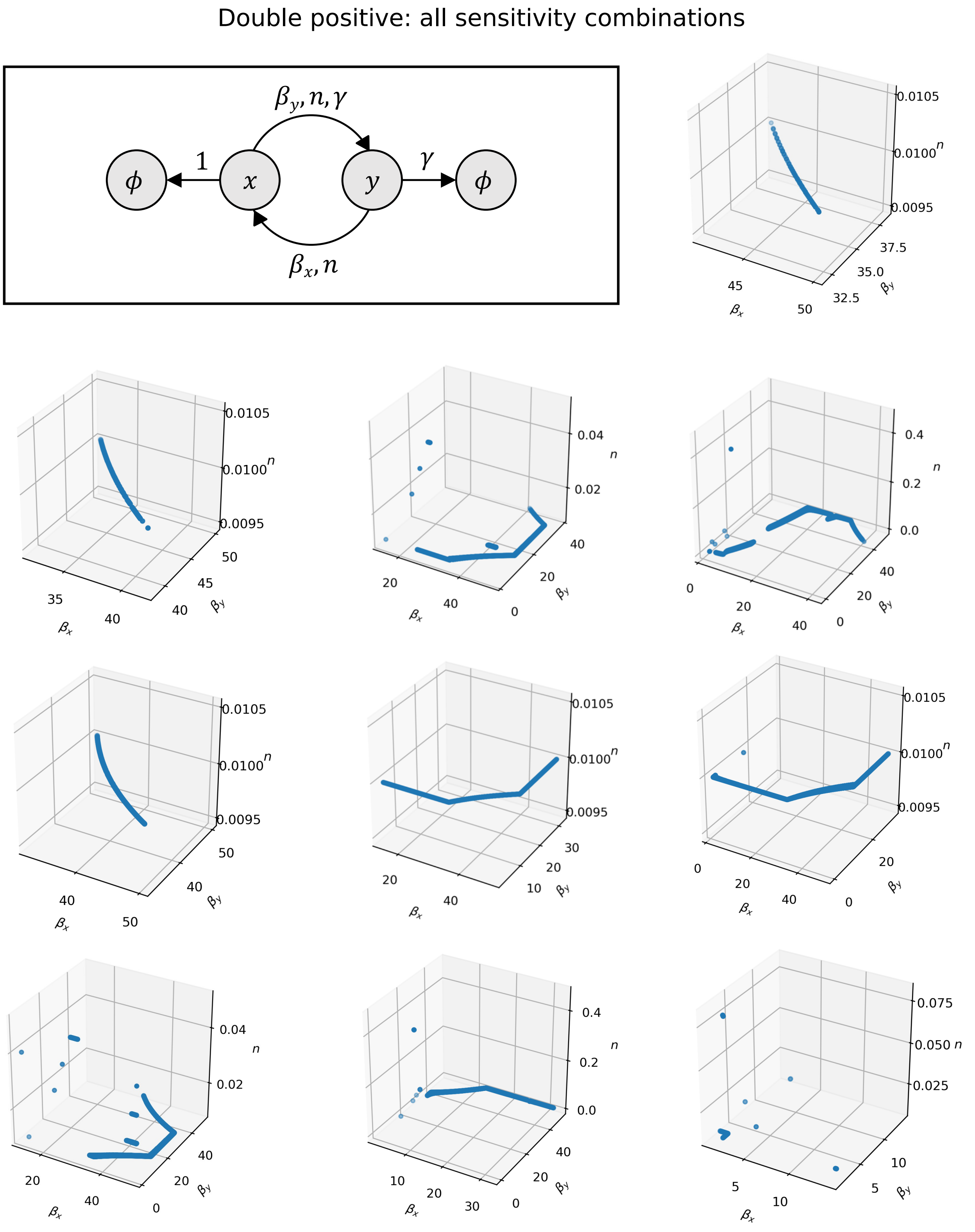


Figure S2: Parameters corresponding to each Pareto front in Figure S1. Each panel matches to the panel in Figure S1.

Positive-negative: all sensitivity combinations

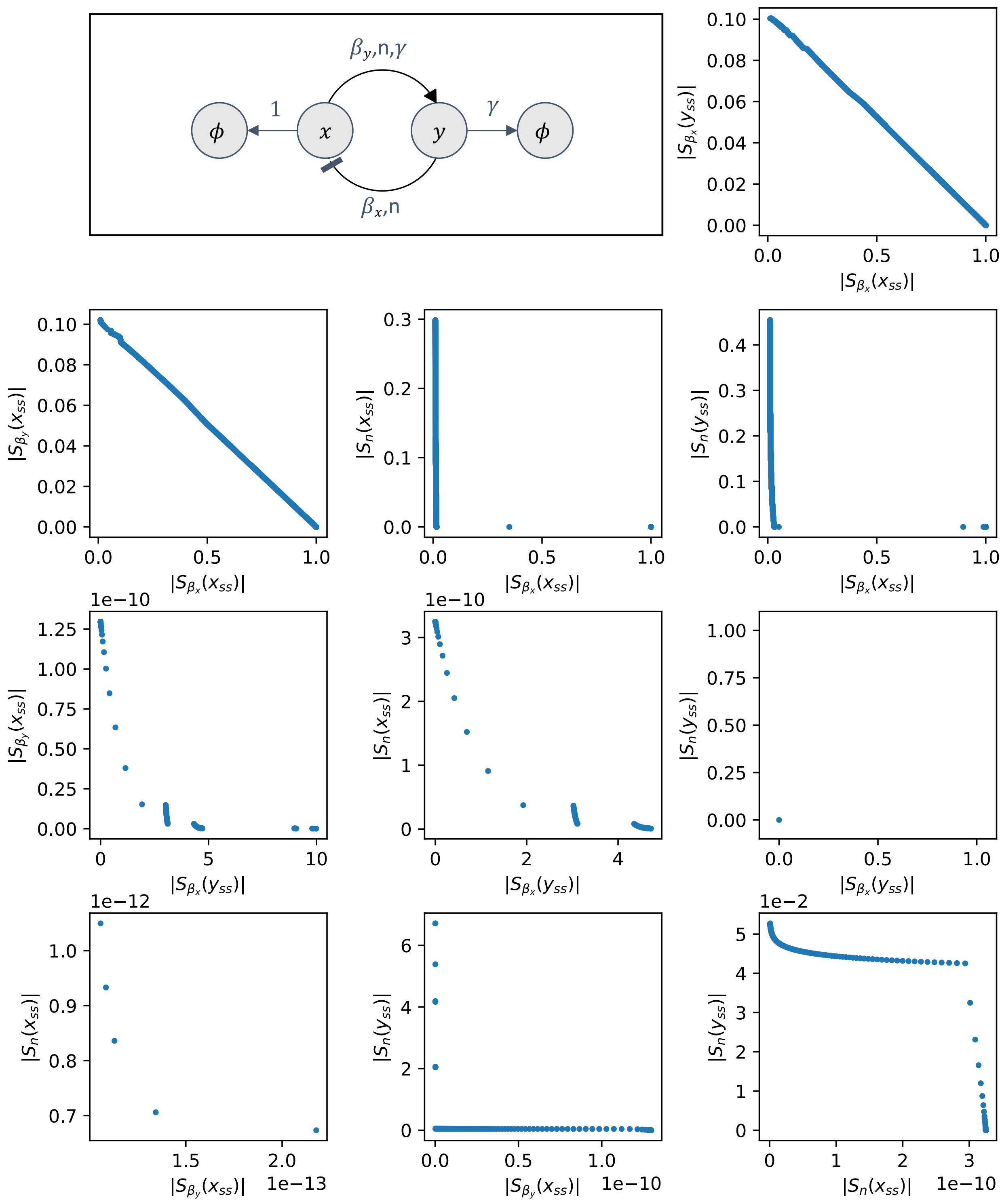


Figure S3: Pareto fronts obtained from the simultaneous minimization of all pairs of sensitivity combinations for the positive-negative feedback loop.

Positive-negative: all sensitivity combinations

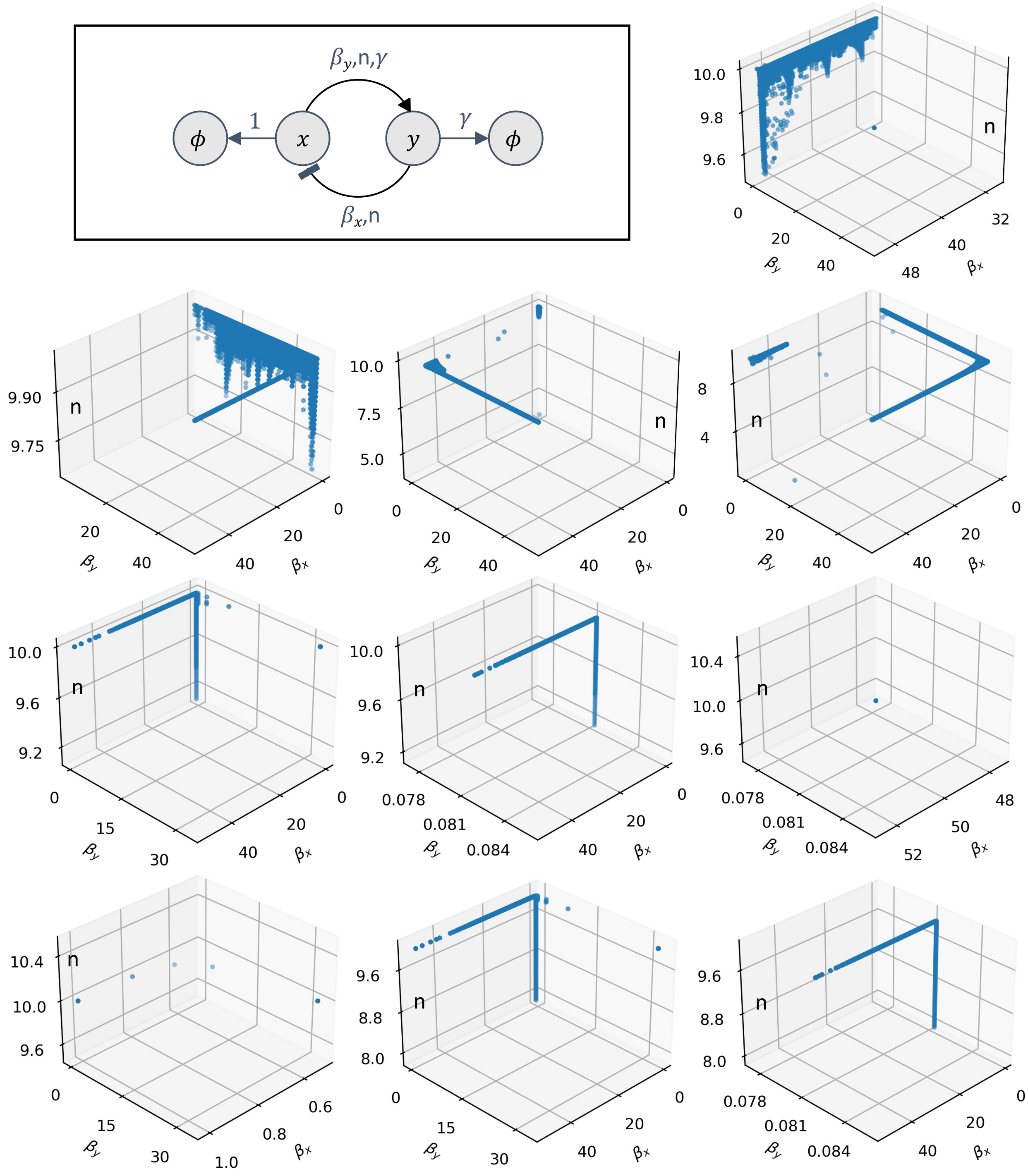


Figure S4: Parameters corresponding to each Pareto front in Figure S3. Each panel matches to the panel in Figure S3.

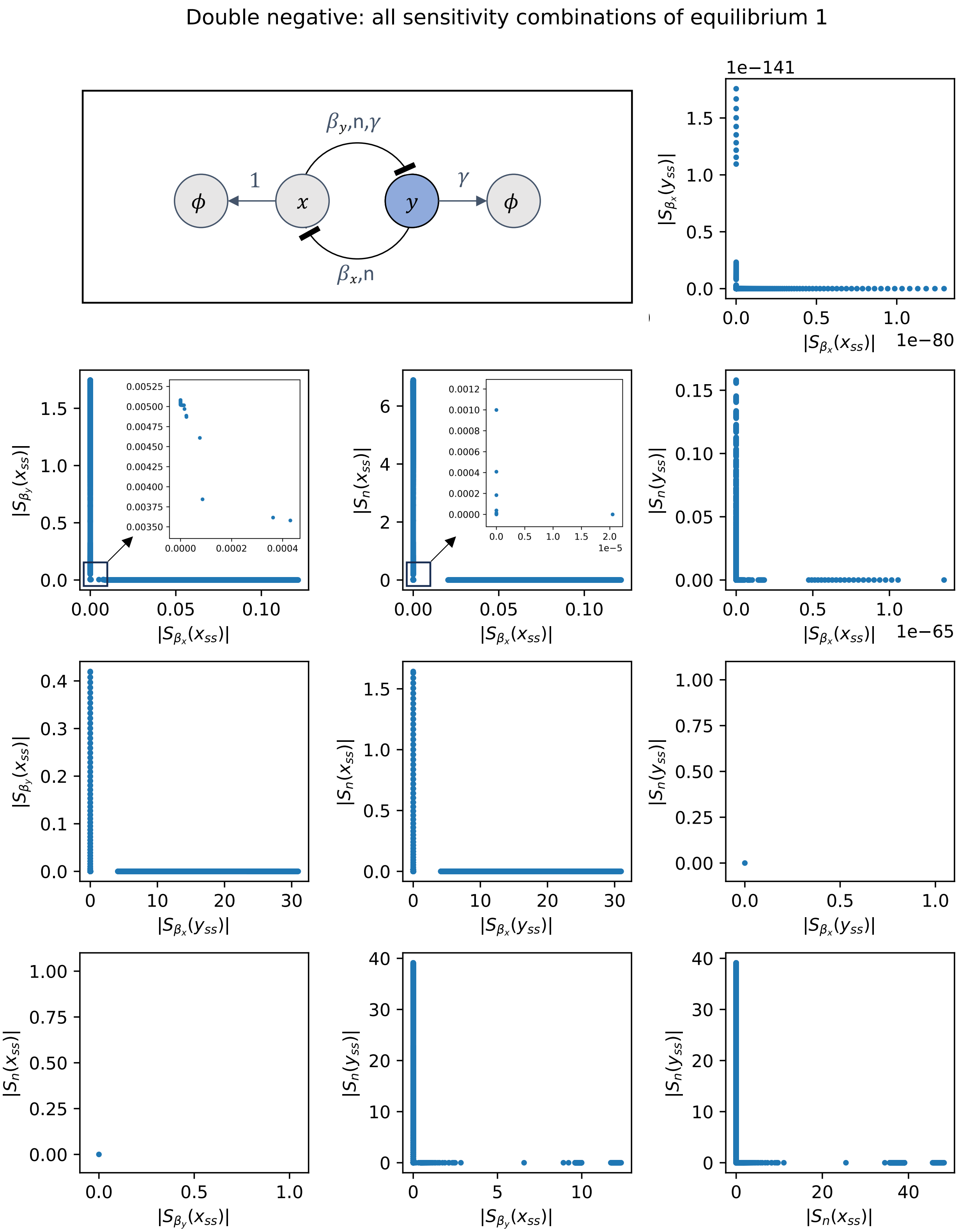


Figure S5: Pareto fronts obtained from the simultaneous minimization of all pairs of sensitivity combinations for the steady state associated with species $y$ overtaking species $x$ in the toggle system (double negative feedback).

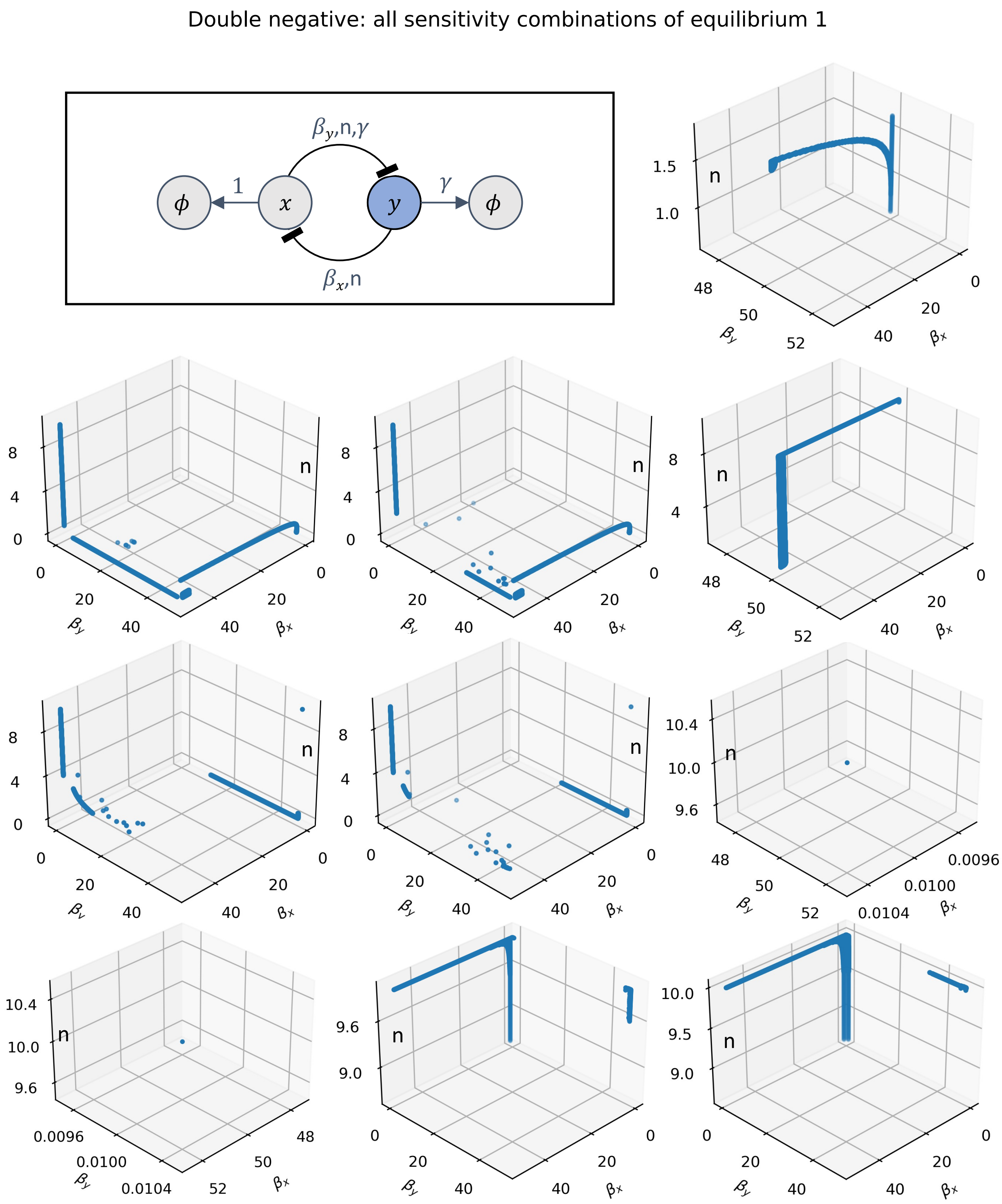


Figure S6: Parameters corresponding to each Pareto front in Figure S5. Each panel matches to the panel in Figure S5.

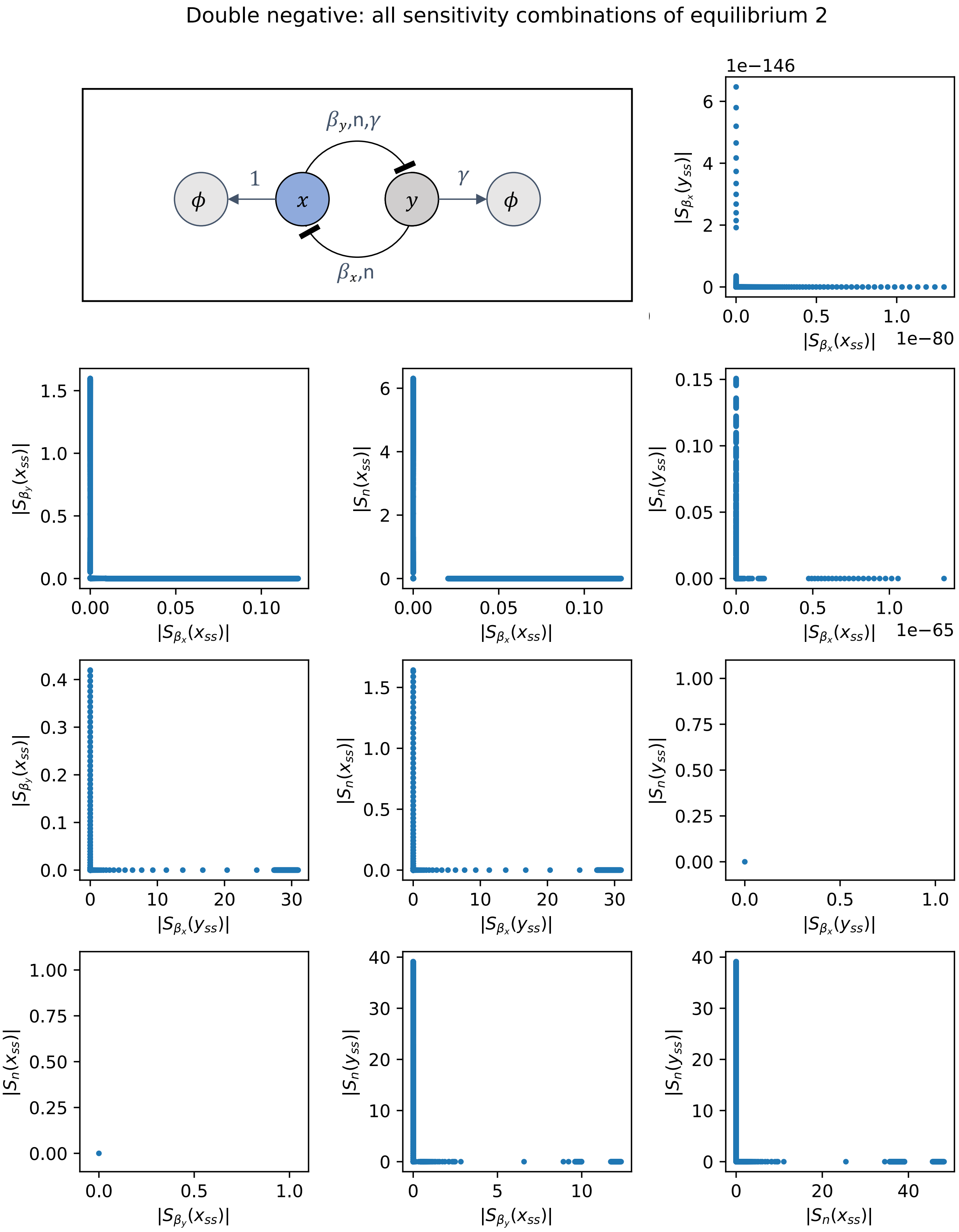


Figure S7: Pareto fronts obtained from the simultaneous minimization of all pairs of sensitivity combinations for the steady state associated with species $x$ overtaking species $y$ in the toggle system (double negative feedback).

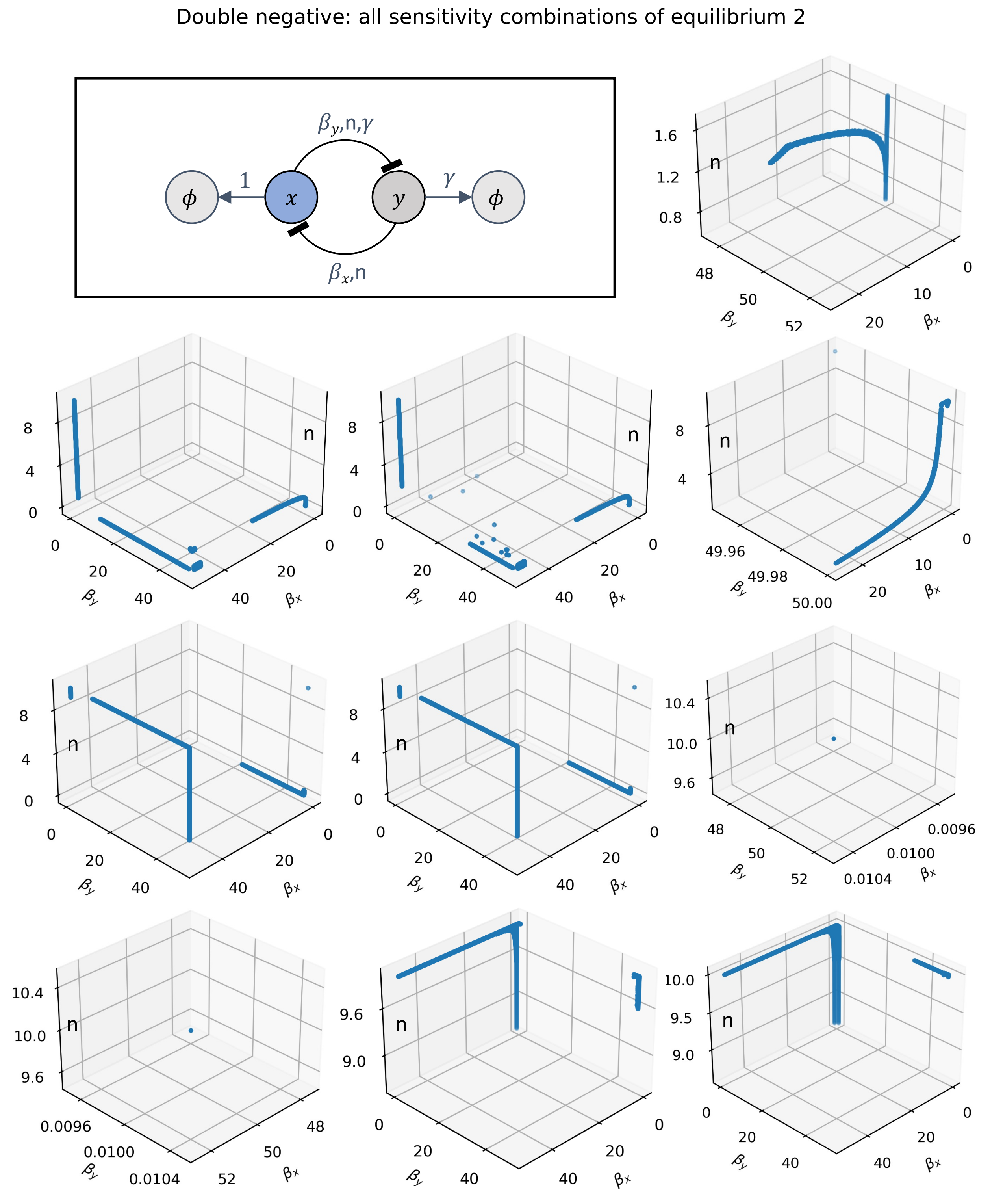


Figure S8: Parameters corresponding to each Pareto front in Figure S7. Each panel matches to the panel in Figure S7.

For double positive feedback, positive-negative feedback, and double negative feedback architectures, we present representative examples illustrating how leakiness enhances the robustness of a single pair of sensitivity functions in each circuit. The complete set of figures for these two-species circuits, comprising 75 plots, is available on our GitHub repository, as including all of them exceeds the scope of this supplementary document.

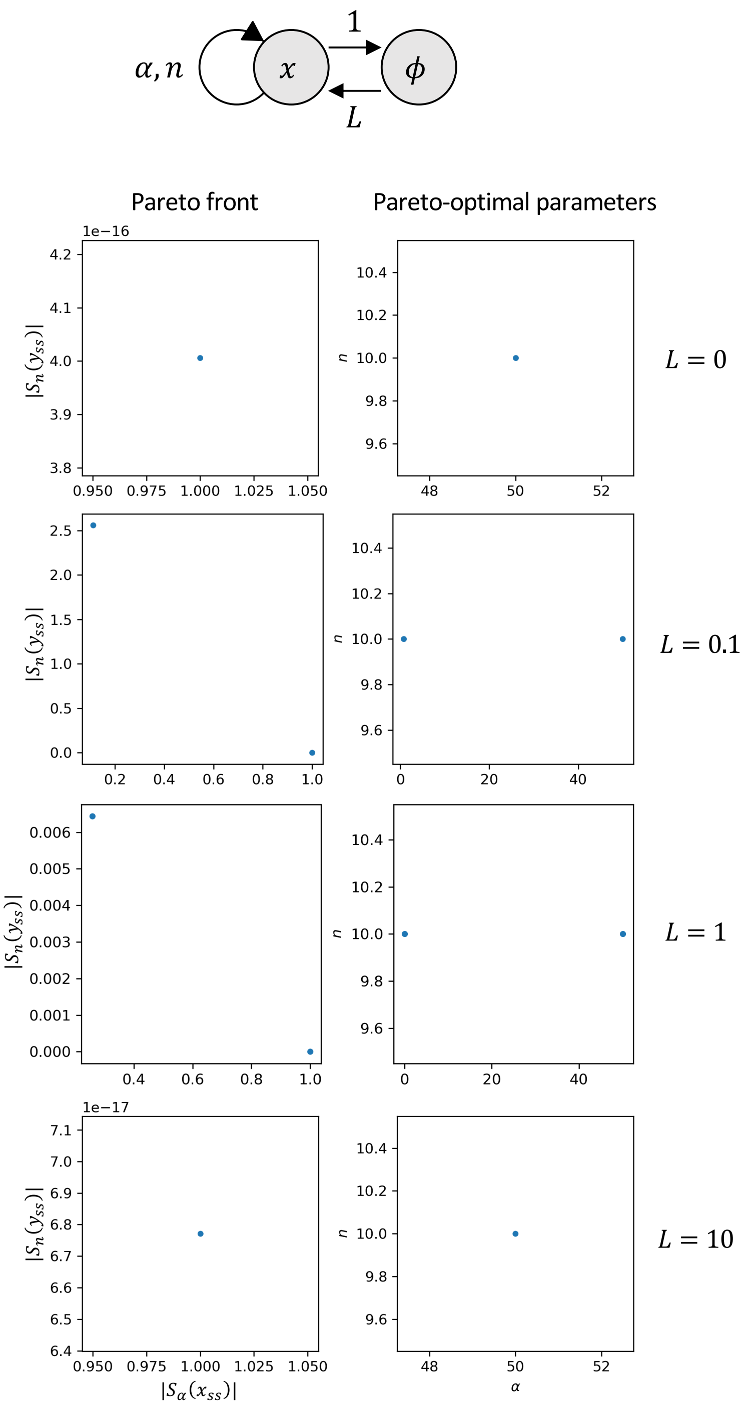


Figure S9: **Positive autoregulation retains maximal robustness despite promoter leakiness.** Initially, at L = 0 , a single Pareto point is observed at $(S_\alpha(x_{ss}), S_n(x_{ss})) = (1, 0)$. This point remains consistent across all subsequent values of $L$ . It is worth noting the appearance of stray Pareto points at $L = 0.1$ and $L = 1$ , which are numerical artifacts and can be disregarded. Additional sampling at these intermediate $L$ values would be required to accurately recover the true Pareto front. For consistency, however, the same parameter sampling resolution of $5000 \times 5000$ is maintained across all plots.

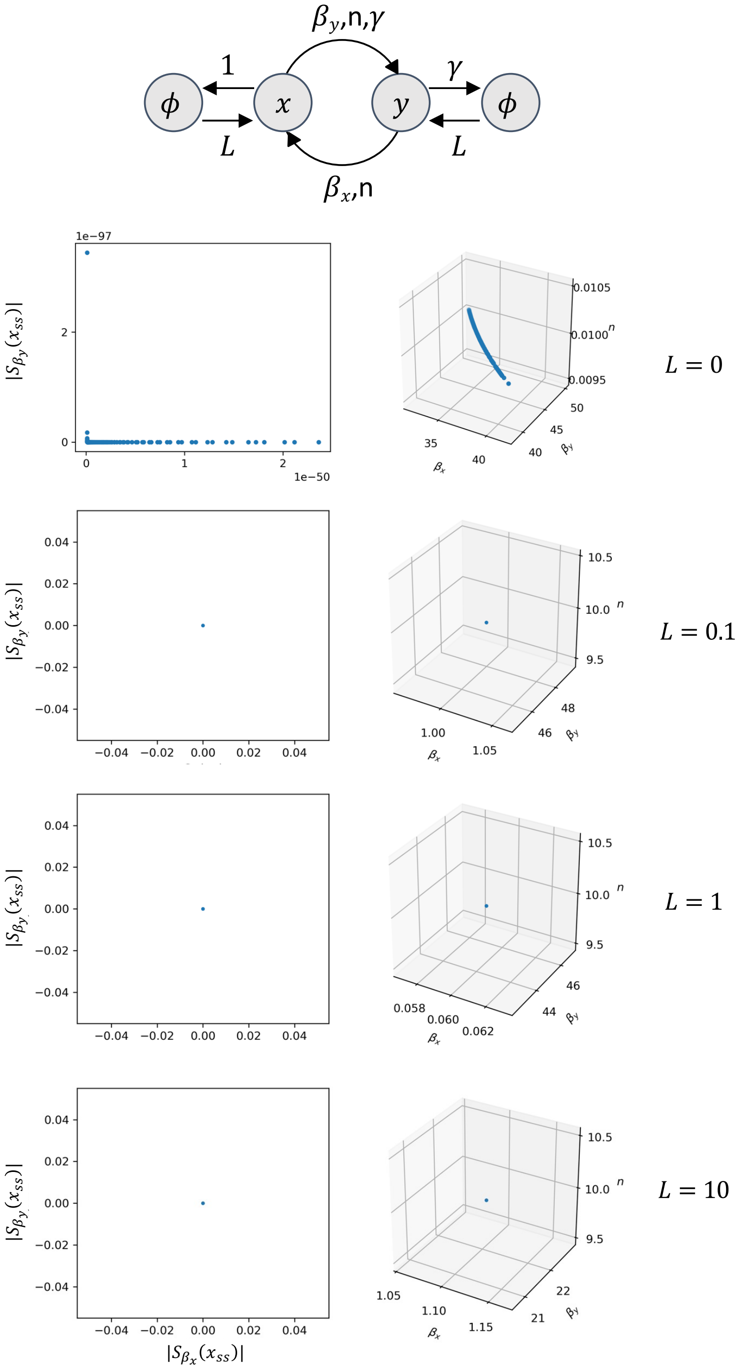


Figure S10: **Double-positive feedback retains robustness with increast promoter leakiness.** No tradeoff changes are observed for double-positive feedback as leakiness increases.

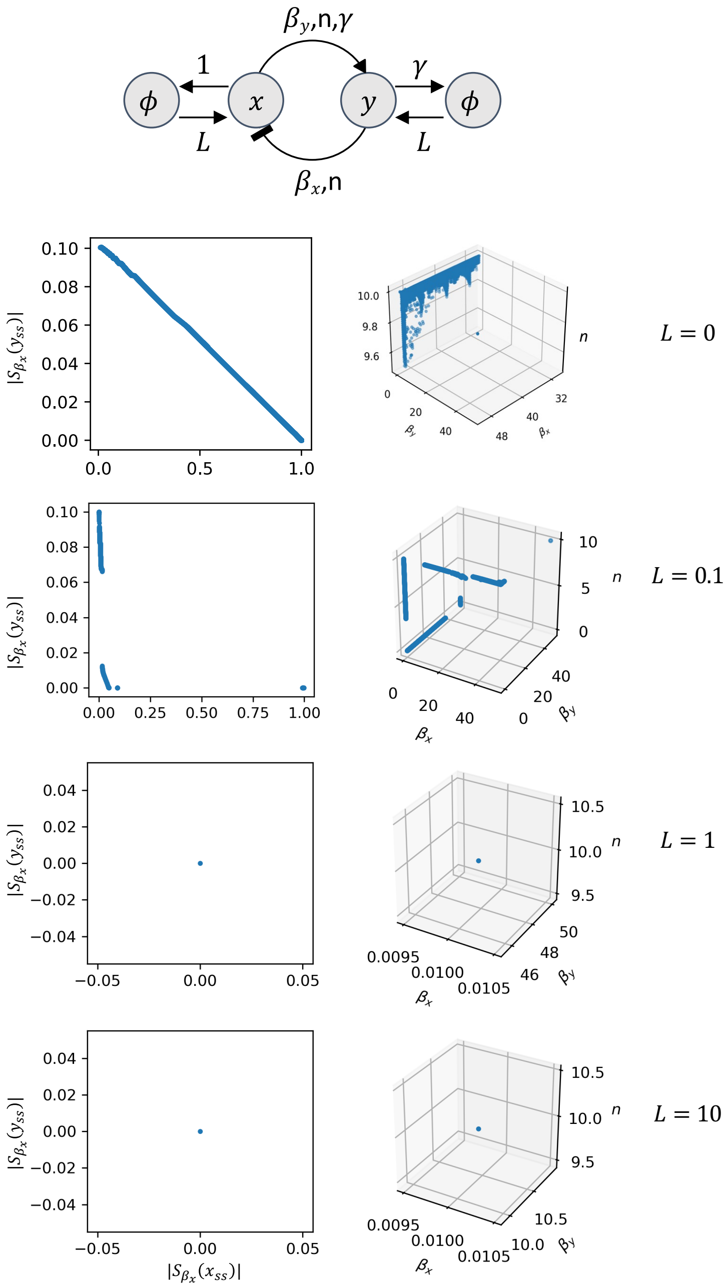


Figure S11: **Positive-negative feedback demonstrates enhanced robustness with increasing promoter leakiness.** Initially, at $L = 0$, the circuit exhibits a tradeoff. At $L = 0.1$, the Pareto front shifts closer to the origin, indicating a reduction in tradeoff. By $L = 1$ and beyond, the tradeoff disappears entirely, and the system converges to a single maximally robust configuration.

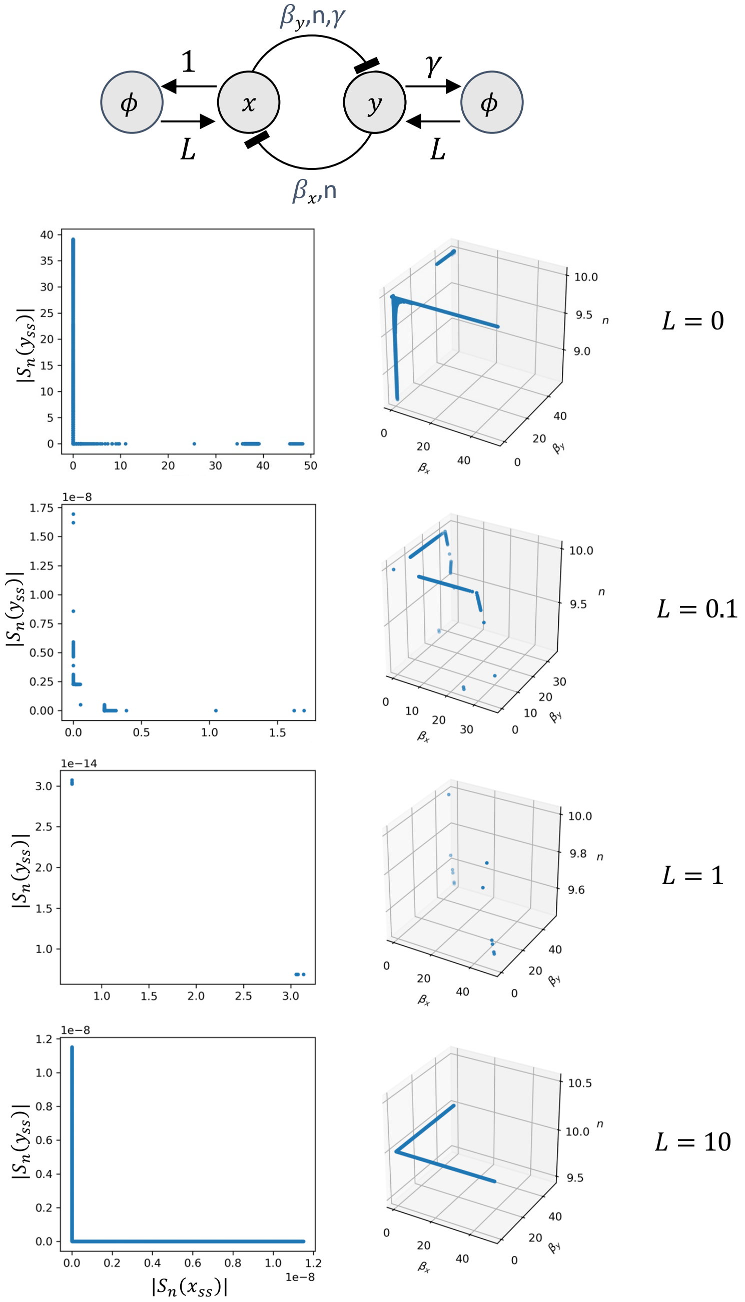


Figure S12: **Double-negative feedback exhibits increased robustness with greater promoter leakiness.** At a parameter sampling resolution of 1000 $\times$1000 $\times$ 1000 , higher levels of leakiness shift the detected Pareto front closer to $(0, 0)$ — the expected Pareto front. This indicates that the sampled parameter space becomes more concentrated and aligned near $(0, 0)$, spanning a more robust range of sensitivities for the same set of sampled parameters.
